# Domestication of different varieties in the cheese-making fungus *Geotrichum candidum*

**DOI:** 10.1101/2022.05.17.492043

**Authors:** Bastien Bennetot, Jean-Philippe Vernadet, Vincent Perkins, Sophie Hautefeuille, Ricardo C. Rodríguez de la Vega, Samuel O’Donnell, Alodie Snirc, Cécile Grondin, Marie-Hélène Lessard, Anne-Claire Peron, Steve Labrie, Sophie Landaud, Tatiana Giraud, Jeanne Ropars

## Abstract

Domestication is an excellent model for studying adaptation processes, involving recent adaptation and diversification, convergence following adaptation to similar conditions, as well as degeneration of unused functions. *Geotrichum candidum* is a fungus used for cheese making and is also found in other environments such as soil and plants. By analyzing whole-genome data from 98 strains, we found that all strains isolated from cheese formed a monophyletic clade. Within the cheese clade, we identified three genetically differentiated populations and we detected footprints of recombination and admixture. The genetic diversity in the cheese clade was similar as that in the wild clade, suggesting the lack of strong bottlenecks. Commercial starter strains were scattered across the cheese clade, thus not constituting a single clonal lineage. The cheese populations were phenotypically differentiated from other populations, with a slower growth on all media, even cheese, a prominent production of typical cheese volatiles and a lower proteolytic activity. One of the cheese clusters encompassed all soft goat cheese strains, suggesting an effect of cheese-making practices on differentiation. Another of the cheese populations seemed to represent a more advanced stage of domestication, with stronger phenotypic differentiation from the wild clade, harboring much lower genetic diversity, and phenotypes more typical of cheese fungi, with denser and fluffier colonies and a greater ability of excluding cheese spoiler fungi. Cheese populations lacked two beta lactamase-like genes present in the wild clade, involved in xenobiotic clearance, and displayed higher contents of transposable elements, likely due to relaxed selection. Our findings suggest the existence of genuine domestication in *G. candidum*, which led to diversification into different varieties with contrasted phenotypes. Some of the traits acquired by cheese strains indicate convergence with other, distantly related fungi used for cheese maturation.

## Introduction

Understanding how populations adapt to their environment is a key question in evolutionary biology. Domestication, the change in the genetic and phenotypic make-up of populations under human artificial selection, is an excellent model for studying adaptation processes, as it involves recent adaptation events under strong selection on known traits, rapid diversification and reduced gene flow between wild and domesticated populations. Numerous studies have documented the specific traits acquired in domesticated animals (dog, horse, pig, cow) ^1–4^ and plants (cabbage, corn, wheat) ^5–7^, as well as their genetic differentiation from wild populations and their adaptive genomic changes. For example, some domesticated animals (e.g. dog, horse and cattle) have been selected for similar traits such as coat color, size, rapidity and docility ^8–10^. In plants too, similar traits have been selected in different lineages, such as bigger grains with more nutrients and lack of dormancy ^5,7,11,12^. On the other hand, functions essential in wild environments but unused in anthropic environments have often degenerated due to relaxed selection, for example, reductions in defense mechanisms in plants ^5,11^. Domestication also often leads to strong reduction in genetic diversity due to bottlenecks in animals (e.g. dog) ^13^ and annual plants (e.g. rice) ^14^. Humans have domesticated several fungi for the fermentation of foods (e.g. beer, bread, wine, coffee, cacao, dry-cured meat and cheese), to produce secondary metabolites used in pharmaceutics (e.g. penicillin), or for their nutritional and gustatory values (e.g. button and shiitake mushrooms) ^15^. Fungi are excellent models for studying evolution and adaptation in eukaryotes, given their many experimental assets ^16^: fungi have relatively small genomes, many are easy to culture in laboratory conditions, and spores can survive long periods in the freezer. However, despite their economic and industrial importance, and their utility as biological models for studying adaptive divergence, the domestication in fungi has yet been less studied than in plants or animals. An exception is the budding yeast *Saccharomyces cerevisiae* used in the production of beer, wine, sake, bread, coffee and cacao ^17–29^, and to a lesser extent the filamentous fungus *Aspergillus oryzae* used to ferment soy and rice products in Asia ^30–32^ and the *Penicillium* species used for cheese ripening, e.g. *P. camemberti* for soft cheeses ^33^, and *P. roqueforti* for blue cheeses ^34–36^. Phenotypic traits beneficial for food production have been acquired in fungal domesticated populations, being different from wild populations. The domestication having led to *P. camemberti* occurred in several steps, with the successive differentiation of several lineages displaying decreasing diversity and increasing beneficial traits for cheese maturation, from the wild *P. fuscoglaucum*, to *P. biforme* and then the two clonal *P. camemberti* varieties, *caseifulvum* and *camemberti* ^33^. Domesticated populations of fermented food microorganisms can for example better assimilate the carbon sources present in the anthropic environment, *e.g.* lactose for *Penicillium* cheese fungi ^36^ and maltose for *S. cerevisiae* sourdough strains ^37^. Furthermore, volatile organic compounds crucial for cheese flavor are more appealing in cheese populations compared to wild populations in *P. roqueforti* ^38^.

The genomic processes involved in adaptation to human-made environments in domesticated fungi include gene family expansion for specific metabolism pathways, gene gain, inter-specific hybridization, introgression and horizontal gene transfer ^22,26,32,34,36, 39–43^. Domesticated fungi also have lost parts of genes no longer useful in the food environment; for example a cheese *P. roqueforti* population and *P. camemberti* var. *caseifulvum* are no longer able to produce some of their toxins due to deletions in the corresponding genes ^33,44^. Such losses are likely due to relaxed selection in terms of competition ability in cheese, in which desired fungi are often inoculated in large quantities compared to possible competitors. Bottlenecks (leading to genetic drift) and degeneration have also been documented in domesticated fungi, with reduced fertility and genetic diversity in the cheese fungi *P. roqueforti* and *P. camemberti* ^33,35^, likely due to an accumulation of deleterious mutations because of drift.

While several cheese-making fungi have been studied recently, it is important to add study cases in additional lineages, as it allows addressing the question of whether adaptation to a similar medium leads to convergent traits. In the case of cheese-making fungi, one can expect phenotypic convergence, for example for more or less rapid growth on cheese, higher proteolysis and lipolysis abilities, higher competitive abilities and greater production of positive volatile compounds ^33^, as these traits may have been selected similarly in the different fungi used for cheese making. *Geotrichum candidum* (teleomorph *Galactomyces candidus*) is a dimorphic fungus (i.e., able to grow as a yeast or a mycelial form), commonly used for cheese-making, but also thriving in other environments such as soil, plants and fruits. *Geotrichum candidum* is naturally present in raw milk and is also often added as a starter culture for the production of semi-hard, mold-ripened, smeared soft cheeses, fresh goat and ewe cheeses. Analyses based on genetic markers have revealed genetic differentiation between cheese and wild strains ^46–49^. Phenotypic diversity within *G. candidum* has been reported in terms of carbon and nitrogen assimilation, lipolysis and proteolysis ^48,50^. However, it has not been tested whether *G. candidum* cheese populations have evolved specific traits that could be beneficial for cheese making.

By analyzing the genomes of 98 strains isolated from different kinds of cheeses and other environments, we confirmed the genetic differentiation between cheese and wild strains and the occurrence of recombination. Within the cheese clade, we reveal the existence of three varieties, i.e., three genetic clusters with contrasted traits and levels of diversity. One of the cheese clusters encompassed all goat soft cheese strains, suggesting an effect of cheese-making practices on differentiation. Commercial strains did not belong to a single clonal lineage, but were instead present in all cheese clusters, some corresponding to admixed strains. We found phenotypic differentiation between cheese and wild populations, and between cheese populations, in terms of growth, proteolysis and volatile compounds. The cheese clade lacked two tandem beta lactamase-like genes, present in wild strains, and involved in xenobiotic clearance, and contained more repetitive elements than the wild clade. Altogether our findings suggest the existence of genuine domestication in *G. candidum,* with both genetic and phenotypic differentiation of cheese strains from their wild counterparts, and a stronger domestication syndrome in one of the cheese clusters. We also show convergence in some traits with domesticated *Penicillium* cheese fungi, such as a denser and fluffier mycelium, at the expense of radial growth, and the production of typical cheese volatiles.

## Results

### Genetic differentiation between wild and cheese strains in *Geotrichum candidum*

We collected and sequenced the genomes of 88 *G. candidum* strains with Illumina technology and included in our analyses ten available genomes (Illumina and PacBio) ^48^. Our dataset included 61 strains isolated from different kinds of cheeses (semi-hard, mold-ripened, smeared soft and fresh soft goat cheeses), 16 industrial strains used for cheese-making, seven strains from dairy products, four strains from other food substrates (e.g., sausage or vegetables) and all the 10 wild strains available in public collections worldwide (isolated for exemple from soil or plan; Table S1). We identified 699,755 SNPs across the 98 strains by mapping against the CLIB 918 reference genome (cheese strain, NCBI accession: PRJEB5752). All the 98 *G. candidum* strains analysed were haploid.

The maximum likelihood tree, principal component analysis (PCA) and neighbor-net (SplitsTree) analyses all identified the same three clades (Figure 1A), with one containing mostly wild strains (corresponding to the GeoC group identified previously based on genetic markers) ^48^, one composed of strains of varying origins (i.e. dairy products and other environments, corresponding to the group previously named GeoB) and one containing mostly cheese and dairy strains (previously named GeoA). The larger sampling and the genome sequencing of the present study further revealed genetic subdivision in the cheese clade, with three clearly differentiated populations and several admixed strains (Figure 1B). We performed an admixture analysis, assigning strains to *K* populations, with *K* ranging from two to ten. At *K*=3, the cheese, mixed-origin and wild clades were separated (Figure S1). At *K*=5, the cheese clade was divided into three genetic clusters, corresponding to monophyletic groups in the maximum likelihood tree and well-separated genetic clusters in the PCA and the neighbor-net (Figure 1 B, D). The second order rate of change of the likelihood (deltaK) peaked at *K*=6. The additional population distinguished at *K*=6 compared to K=5 however encompassed only two strains, that were not much differentiated from others in the splitsTree (MUCL 14462 and CBS 9194, isolated from non-cheese substrates and clustering with the wild population at K=5; Figure 1B). We therefore chose to consider only the five largest populations in the following, as we could not run phenotypic tests on a population with only two strains.

**Figure 1:**
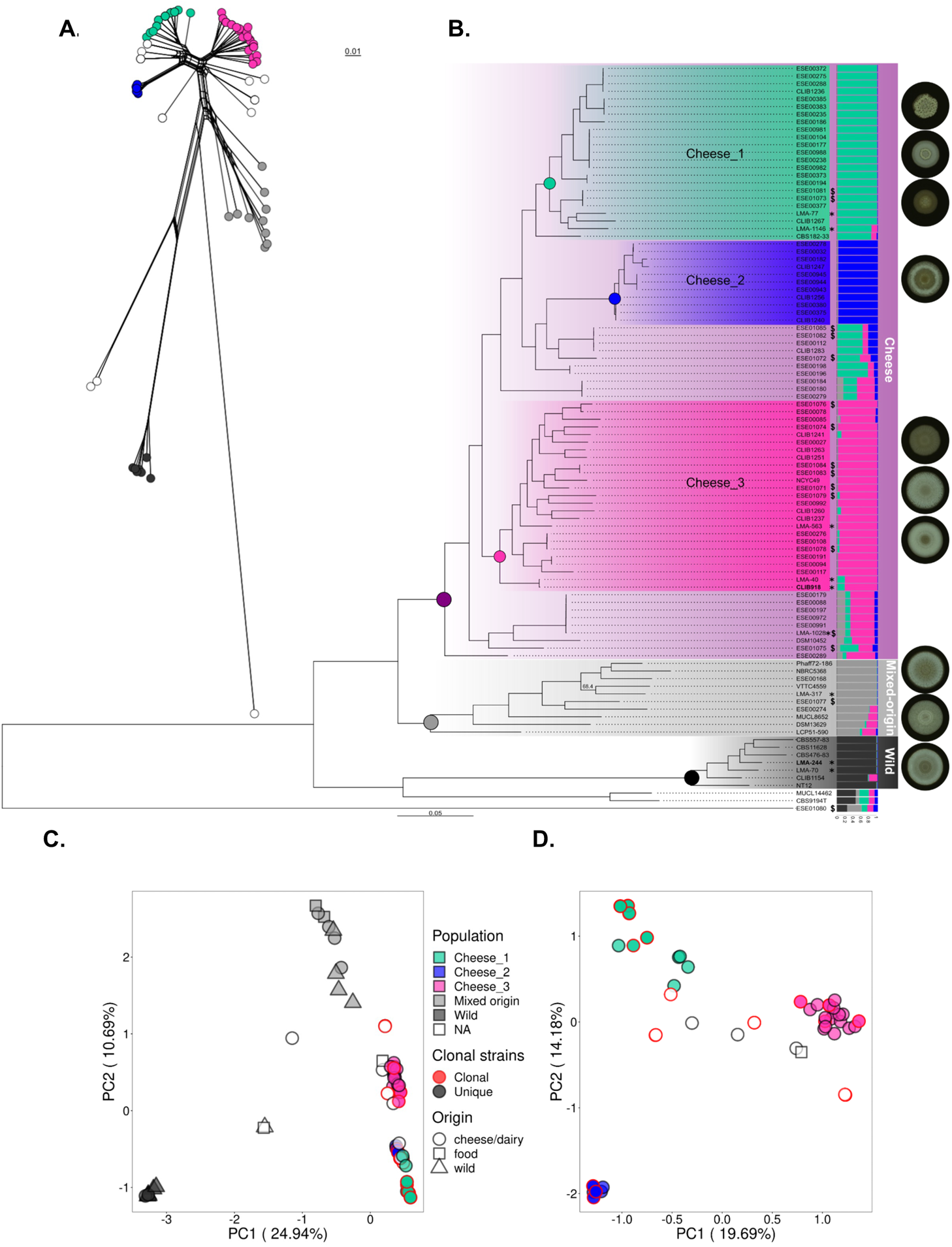
Phylogenetic relationships and population structure of 98 strains of *Geotrichum candidum*, based on whole-genome data. (A) Neighbor-net analysis based on a single nucleotide polymorphism (SNP) distance matrix. The scale bar represents 0.01 substitutions per site for branch lengths. (B) Genetic relationships among strains and population structure in *G. candidum* based on 699,755 SNPs. a) Maximum likelihood tree showing phylogenetic relationships among the 98 isolates used in this study. All nodes are supported by bootstrap support >98% (bootstrap analysis with 1000 resampled datasets). The scale bar represents 0.05 substitutions per site for branch lengths. We used the midpoint rooting method to root the tree. The “$” symbol pinpoints commercial starter strains and “*” the PacBio sequences. Genomes used as references are written in bold. b) Population subdivision inferred for *K* = 5. Colored bars represent the coefficients of membership in the five gene pools based on SNP data. (C) Principal component analysis (PCA) based on 699,755 SNPs and 98 strains. Genetic clusters are represented by the same colors on all panels: light blue for Cheese_1, dark blue for Cheese_2, pink for Cheese_3, light grey for the mixed-origin population and dark grey for the wild population. The symbols represent the environment from which strains were sampled: circle for cheese/dairy, square for food and triangle for wild environment. Symbols are circled in red when multiple strains are overlapping due to clonal lineages (with a threshold set at <1,200 SNP for defining clonal strains). (D) PCA based on the 323,385 SNPs when the dataset was restricted to the 78 strains from the cheese clade.

Some cheese strains could not be assigned to any genetic cluster with the admixture analysis and were placed on the PCA between the three well-delimited cheese genetic clusters (Figure 1C-D), suggesting that they resulted from admixture events. To test this hypothesis, we investigated whether these strains had mosaic genomes, with different genomic regions assigned to distinct clusters. We calculated pairwise identity between unassigned strains and the other strains, computing mean identity to the different genetic clusters along scaffolds using sliding windows. For all unassigned strains in the cheese clade, we observed shifts in identity values along scaffolds, confirming that these strains are the results of admixture between clusters (Figure S2). In contrast, the three unassigned strains outside of the cheese clade did not show changes in similarity level to the different clusters along their genome; these strains may belong to yet additional genetic clusters that could not be distinguished by the analyses because too few strains belonged to these clusters in the sampling (Figure 1B). We tested for an over-representation in the three cheese clusters of the five types of cheeses from which strains were sampled, i.e., soft, soft natural rind, pressed uncooked, pressed cooked and blue cheeses (Table S1). We found an over-representation only in the cheese_1 population (chi2 = 36.6 ; df = 12 ; p < 0.001), which encompassed all strains isolated from soft natural rind goat cheeses, these kinds of cheeses having a characteristic convoluted aspect. The wild clade was the most differentiated population from all other *G. candidum* populations with *F_ST_* values above 0.70 and *d_xy_* of 1.04E-02 (Table S2). The percentage of private SNPs in the five populations was also high (Table S3). F3 tests based on the number of shared sites (Table S4) supported the genetic differentiation between these populations and the lack of gene flow or migration.

### High nucleotide diversity within cheese populations and footprints of recombination

The overall diversity in the cheese clade (𝜋 = 2.82E-3) was higher than in the wild population (𝜋 = 2.12E-03). Each of the three cheese populations of *G. candidum* had however reduced nucleotide diversities compared to wild and mixed-origin populations (Table S5), by at least a factor of two. The Cheese_2 population showed the lowest genetic diversity (𝜋 = 4.82E-04), by a factor of four compared to the two other cheese populations (Table S5).

*Geotrichum candidum* is a heterothallic fungus, meaning that sexual reproduction can only occur between two haploid cells carrying different mating types. Two mating types have been described in *G. candidum* ^51^: MATA, encoded by a HMG box gene homolog to the MATA2 *Kluyveromyces lactis* allele, present in CLIB 918 (sequence id: HF558448.1), and MATB, encoded by an alphabox gene homolog to the MATα1 *S. cerevisiae* allele, present in the strain CBS 615.84 (sequence id: HF558449.1). In the Cheese_2 population, we found a significant departure from the 1:1 mating-type ratio expected under regular sexual reproduction; all the 12 strains carried the MATB allele, suggesting that this population is at least partly clonal (Table S6). The absence of linkage disequilibrium decay with physical distance between two SNPs (Figure S3), together with the absence of reticulation in the neighbor-net (Figure 1), are also consistent with a lack of recombination in the Cheese_2 population. However, pairwise homology index (PHI) tests, testing with permutations the null hypothesis of no recombination by looking at the genealogical association among adjacent sites, were significant in all the *G. candidum* populations (Table S7); this indicates that recombination did occur at least in a recent past, even in the Cheese_2 population.

We did not detect any accumulation of nonsense or missense mutations in any population compared to silent mutations (Table S8), while degeneration can be expected to be particularly strong in clonally replicated populations as recombination allows more efficient selection. None of the genes that presented nonsense mutation had predicted functions that could be detected as specific either to the wild or cheese environments (Figure S4). This suggests that the absence of sexual reproduction in the Cheese_2 population may be too recent to observe an accumulation of deleterious mutations.

In contrast to the Cheese_2 population, we found both mating-type alleles in balanced proportions in both Cheese_1 and Cheese_3 populations (Table S6) and we observed sharp decays in linkage disequilibrium (LD) with genomic distance, although LD levels remained higher than in the mixed-origin and wild populations (Figure S3). We observed reticulations in the neighbor-net network within populations and, to a lesser extent, between populations (Figure 1A).

As previously mentioned, the 16 commercial starter strains in our *G. candidum* dataset were scattered in the maximum likelihood tree (“$” symbol, Figure 1B.a.) and we detected above footprints of recombination in the Cheese_1 and Cheese_3 populations (Figure 1A, Figure S3). We nevertheless detected a few groups of clonemates, by the lack of branches in the trees and the presence of fewer than 1,200 SNPs between strains (Figure 1; Table S1, clonal group column). As strains within these clonal lineages were isolated from different cheeses and from various French regions, it indicates that these lineages are likely clonally cultivated and sold for cheese making. Some of the commercial starter strains were, in fact, placed within these clonal groups (“$” symbol on Figure 1B). Among the admixed cheese strains, 19 out of 23 were part of clonal lineages.

### Copy number variation: lack of two tandem beta lactamase-like genes in the cheese populations and higher repeat content

Expansions of gene families involved in specific metabolism pathways, of transposable elements and loss of genes no longer required in the new environment can be involved in adaptation to new environments. For example, variations in gene copy number were associated with the adaptation of *S. cerevisiae* to beer making, with duplications of genes involved in maltose and maltotriose metabolism specifically in domesticated beer strains ^22,23,53^. We therefore looked for gene copy-number variation (CNV) that differentiated wild and cheese populations, using 500 bp sliding windows and two reference genomes, belonging to the Cheese_3 and the wild populations, respectively (Table S9). Using the Cheese_3 reference, we found 61 CNV regions (mean length of 1515 bp and 45 non-genic CNVs), encompassing in total 16 genes, half having predicted functions, none being obviously related to cheese adaptation (e.g. methylglyoxal reductase and tRNAs, Table S9). Using the wild genome reference, we found 132 CNV regions (mean length of 1664 bp and 105 non genic CNVs), encompassing 29 genes (seven with unknown functions).

One of these regions, 20 kb long, included only two genes, both matching the Pfam hidden markov model for beta-lactamases; these two genes (g5112 and g5113) were absent from all cheese populations, and were present in most wild strains (except one that had partially the region) and in four strains belonging to the mixed-origin population (Figure 2). The nucleotide identity between the two beta lactamase-like genes was 93%. A third beta lactamase-like gene (g5111) was found immediately next to this CNV region in all *G. candidum* strains, and displayed a nucleotide identity of 87% with the two other beta lactamase-like genes within the CNV. Surrounding these different genes, we found several Tc1/*mariner*, a LINE/Tad1 and other DNA transposons, that may have contributed to the beta-lactamase-like gene deletion (Figure 2). Fungal beta lactamase-like genes are known to contribute to hydrolysis of microbial and plant xenobiotics, and thus may be important in the wild environment to compete with other microorganisms ^54^. If the ancestral state in *G. candidum* is the presence of three beta-lactamase genes, the cheese populations may have lost these two copies of the beta-lactamase genes due to relaxed selection; indeed, these functions may not be useful in the cheese environment if *G. candidum* is inoculated in high quantity compared to potential competitors.

**Figure 2:**
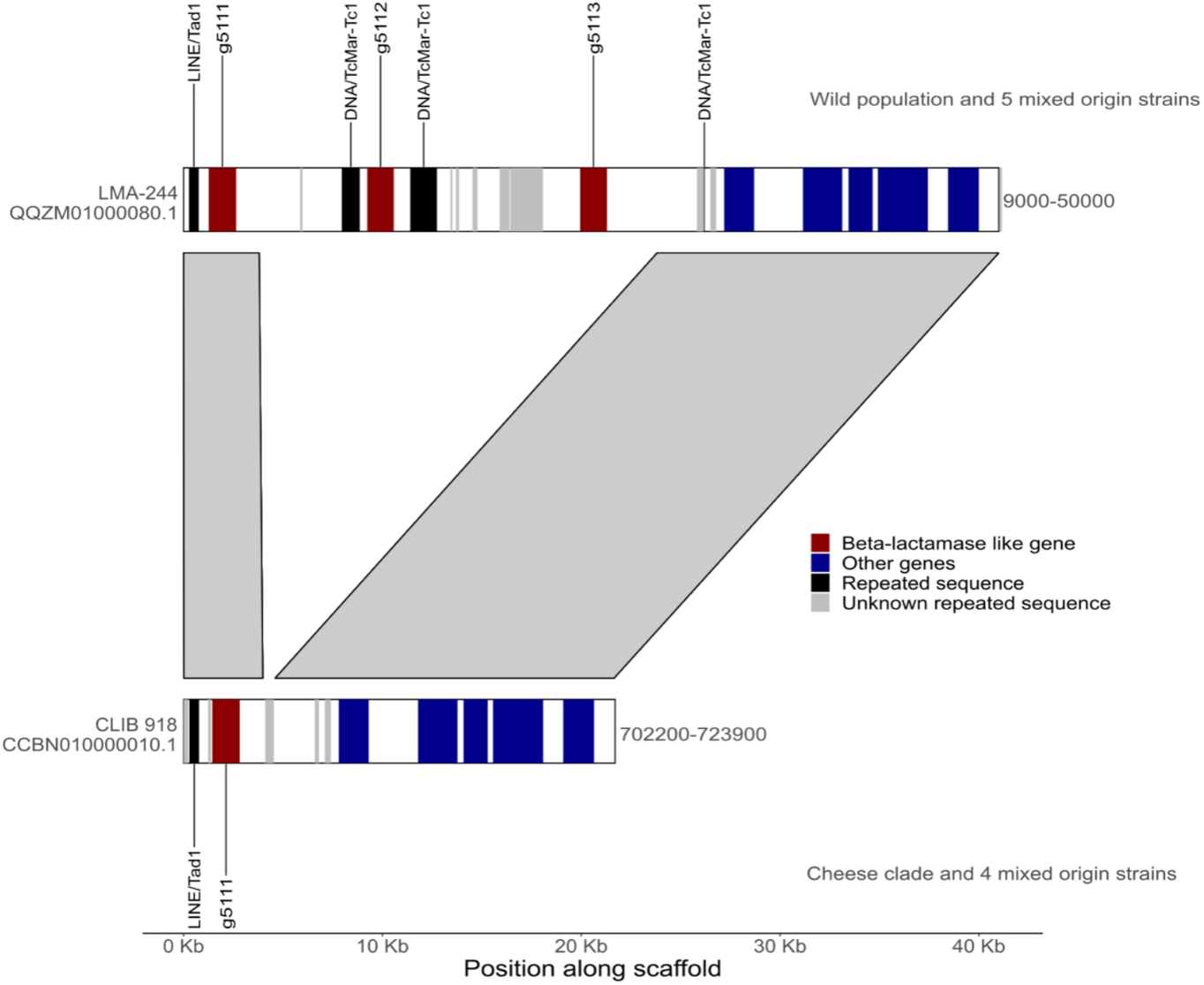
Lack of the beta lactamase-like genes in the cheese clade of *Geotrichum candidum*. Synteny between parts of the scaffold QQZM01000080.1 of the LMA-244 (wild) strain against the scaffold CCBN010000010.1 of the cheese CLIB 918 (Cheese_3) strain. Beta-lactamase-like genes are annotated in red while other genes are displayed in blue. Black triangles indicate positions with repeated sequences. All strains from the cheese clade and five strains from the mixed-origin populations (LMA-317, ESE00274, MUCL8652 and ESE00540) lacked the g5112 and g5113 genes, both encoding for beta-lactamase like.

*De novo* detection of repeats using the wild strain LMA-244 yielded a library containing 107 types of repeated elements (including 15 types of DNA transposons and 11 of retroelements and 3 rolling-circles). We identified 14 types of repeated elements present in at least one other *G. candidum* genome with five times more copies than in the LMA-244 wild strain (this threshold was set based on the fat tail of the distribution; Figure S5). Among these 14 types of repeated elements, several DNA transposons of the Tc1/*mariner* repeat family showed a cheese-clade specific expansion (Table S10, Figure 3). Several unidentified, *Tad1* and *Helitron* repeat types also showed expansions in the cheese clade, alongside a milder expansion in the mixed origin clade. Such higher contents of transposable elements in the cheese clade could result from expansions due to relaxed selection in the cheese clade ^55^.

**Figure 3:**
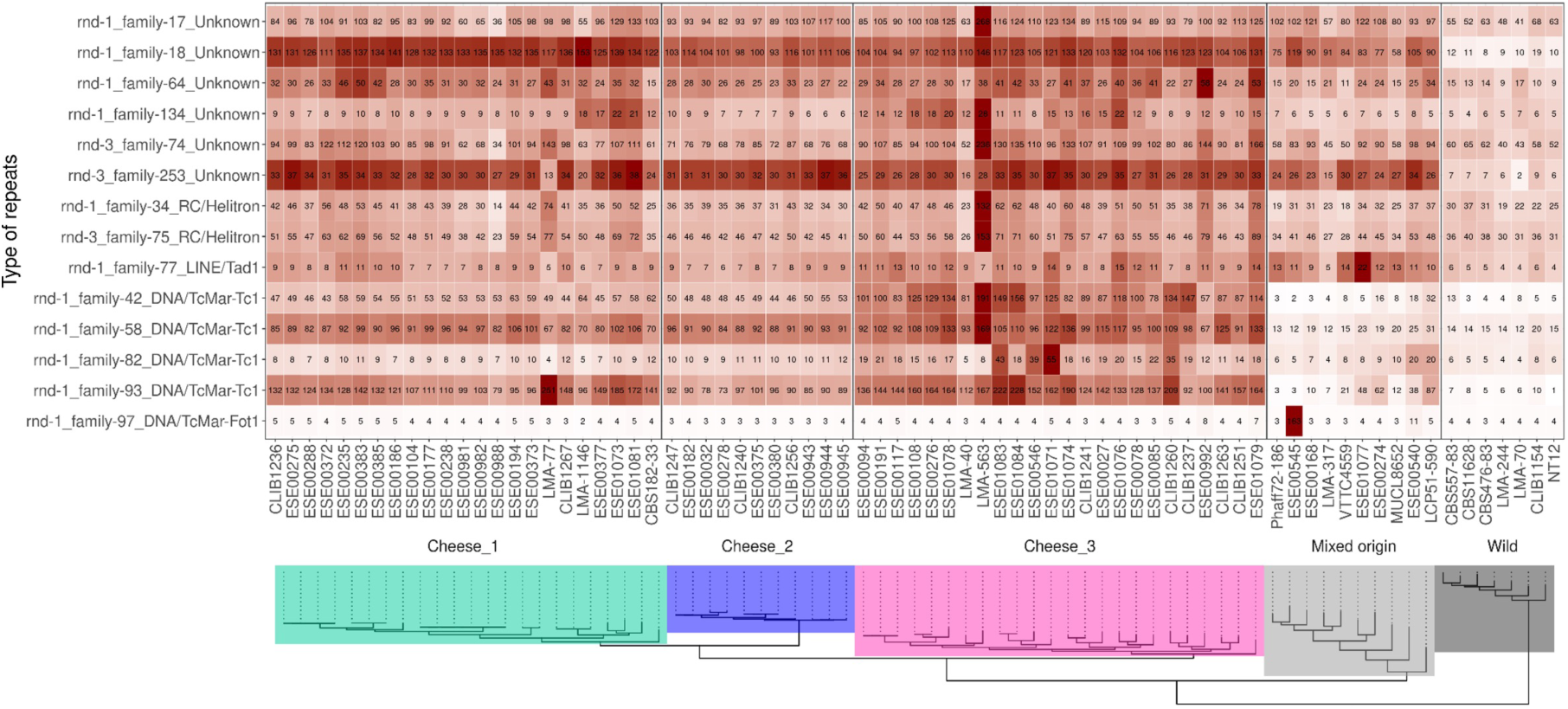
Heatmap of repeats in the cheese populations. The total number of copies is indicated in the center of each cell and by the grey to red color scale. The maximum likelihood (ML) tree from the figure 1 (without admixed strains) is plotted below strain names.

### Genomic footprints of adaptation: genomic islands of differentiation and genes under positive selection

We looked for genomic regions with a greater differentiation or a lower genetic diversity than the genomic background when comparing each of the three cheese populations to the wild population, to detect footprints of divergent selection and recent selective sweeps, respectively. We scanned the whole genome in each cheese cluster using non-overlapping windows and identified the windows within both the 1% highest differentiation with the wild population (*d_xy_*) and the 5% lowest within-population diversity (π). We also performed an analysis using SweeD on each genetic cluster to detect selective sweeps based on the site frequency allelic spectrum, keeping the windows with the 1% highest likelihood. The putative functions of outlier genes are given per cluster and analysis in the Table S11. In the Cheese_1 population, four genes were outliers in all three selection analyses (1% highest likelihood sweeD, 1% highest differentiation and 5% lowest diversity), with one coding for a Ca²⁺-dependent cystein protease (Table S11). This function was also represented in outliers in the other cheese populations, but only shared between the sweeD and d_XY_ analyses. Proteases are important in cheese making as the breakdown of milk caseins greatly contributes to cheese texture and decreases water activity by degrading proteins into molecules with free carboxyl and amino groups ^56^. *Geotrichum candidum* is prevalent during the amino-acid catabolism ripening step of Pelardon fresh cheese ^57^, suggesting that *G. candidum* plays an important role in proteolysis in cheese-making. Among the genes shared between the d_XY_ and sweeD analyses, we also detected functions related to iron uptake, in particular the homolog of the *Schizosaccharomyces pombe* Ctr4 in Cheese_1 and a high-affinity iron permease in Cheese_2. Ctr4 is a copper-sensing transcription factor regulating iron uptake genes in yeasts (Labbé 1999). Iron is limiting in cheese and it may therefore be advantageous for cheese strains to better regulate iron uptake ^58^.

We also searched for genes evolving under positive selection in terms of high rates of non-synonymous substitutions by performing McDonald and Kreitman (MK) tests (Table S12), first by comparing the mixed-origin population (as the closest outgroup) to each cheese population and to the cheese clade as a whole. We detected 25 genes as evolving under positive selection in at least one cheese population (9 for Cheese_1, 18 for Cheese_2, two in Cheese_3 and one in all three cheese populations at once; Table S12). Among them, a metalloendopeptidase evolved under positive selection in all three cheese populations, likely playing a role in casein degradation through cell lysis ^61,62^. A Glucan 1,3-beta-glucosidase was also detected as evolving under positive selection in the Cheese_2 population; this enzyme could be involved in fungal inhibition through fungal cell degradation ^63^. The other genes under positive selection had either no predicted function or putative functions that could not be related to cheese adaptation (Table S12). We also searched for genes with high rates of non-synonymous substitutions by comparing the wild population on the one hand with the mixed-origin and cheese clade on the other hand. We thereby found 23 genes evolving under positive selection (Table S12), with 15 encoding proteins with predicted functions. Among them, a spermidine resistance protein likely plays a role in the yeast-hyphal transition ^64^, *G. candidum* being a dimorphic fungus.

### Phenotypic differentiation between cheese and wild populations

#### Denser mycelial growth and/or faster proteolysis in cheese populations of *Geotrichum candidum*

Cheese fungi selected by humans are expected to display specific traits beneficial for cheese making, such as faster growth in cheese at cave temperature or colonies of attractive aspect or color. In contrast, the ability to grow in harsh conditions, e.g., with little nutrients, may have been lost in cheese strains due to relaxed selection, as often reported for unused traits in human-made environments in domesticated organisms ^36,66,67^.

We therefore measured colony radial growth of 31 strains from the five *G. candidum* populations on different agar media (cheese, rich and poor media) at different temperatures. The wild population grew faster than cheese populations on all media (on a poor medium containing only inorganic salts and a low-concentration carbon source, but also on cheese media) and at all temperatures, with a more pronounced difference at 25°C (Table S13, Figures 4A and S8). This may result from trade-offs with other traits, such as a fluffier mycelium, i.e. more vertical growth at the expense of less radial growth.

To test whether cheese populations had a denser mycelium or had become whiter and/or fluffier, we compared the opacity of populations on cheese agar at cave temperature (10°C), which integrates the brightness and fluffiness of a colony. The Cheese_1 and Cheese_3 populations were not more opaque than wild populations (Figure 4B). The Cheese_2 population had a significantly higher opacity than all other *G. candidum* populations, except the mixed-origin population (post-hoc Tukey test in Table S13). Lipolysis and proteolysis are crucial biochemical processes during cheese ripening, as they influence flavor and texture of the final product. However, too fast proteolysis or lipolysis can lead to degraded products. The wild and mixed-origin populations degraded a significantly higher amount of proteins than the cheese populations while we did not detect any proteolysis in the Cheese_2 population in our experiment (Figure S9; Table S14). All populations had similar lipolysis rates.

**Figure 4:**
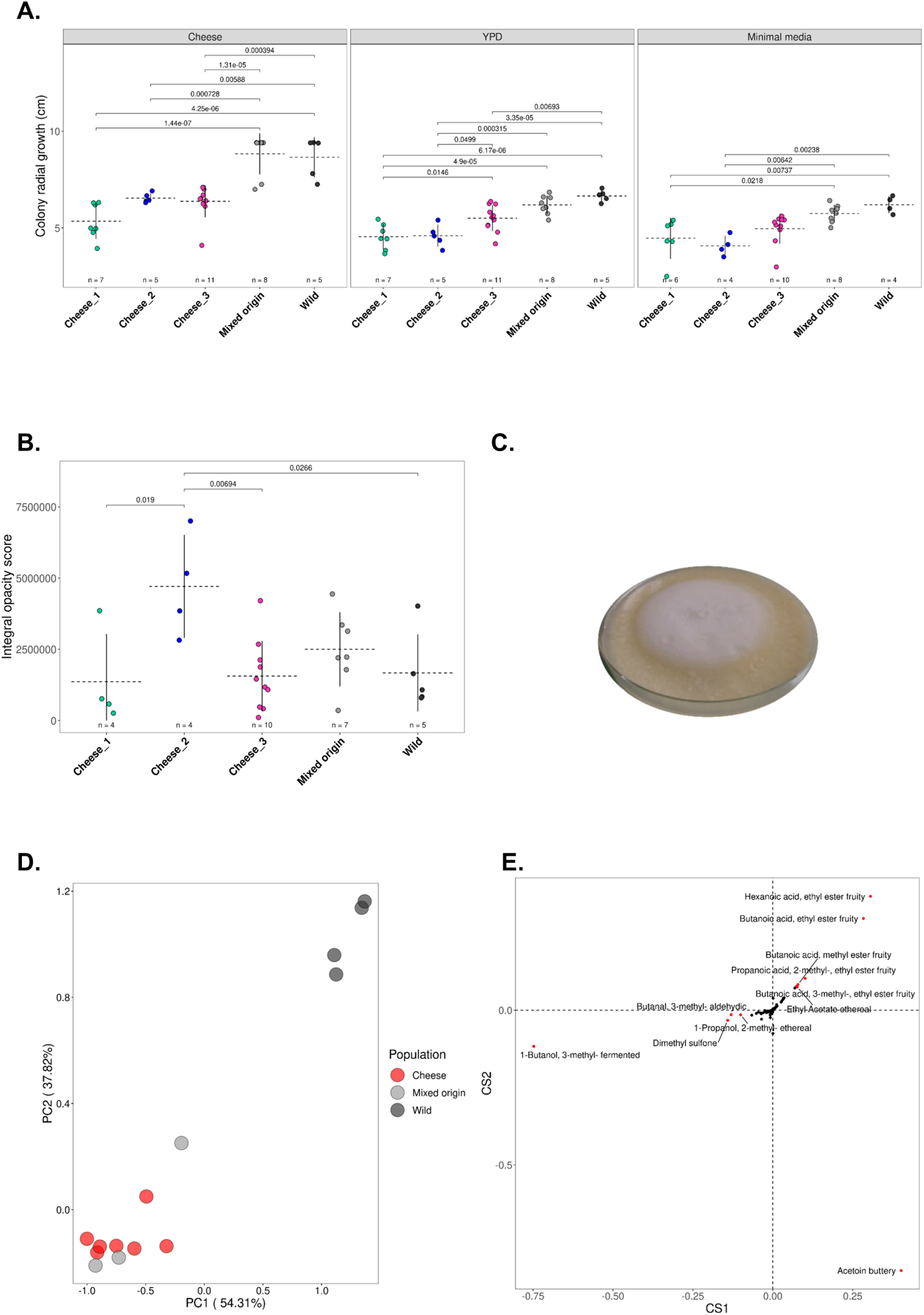
Differences in growth, opacity and volatile compounds between the five populations of *Geotrichum candidum*. Each point represents a strain, horizontal dotted lines and vertical lines represent the mean and the standard deviation of the phenotype in the population, respectively. The number *n* at the bottom of plots indicates the number of strains used per population for measuring the corresponding phenotypes. The pairwise Tukey tests performed to assess whether there were mean differences between populations are indicated between brackets with their p-values. (A) Mean radial growth of the three cheeses, mixed-origin and wild populations on cheese (1% salt), yeast peptone dextrose (YPD) and minimal media at 25°C. (B) Differences in opacity between the three cheese populations, mixed-origin and wild populations on cheese medium (1% salt) at 10°C. Integral opacity is defined as the sum of the brightness values for all the pixels within the fungal colony bounds and measures the whiteness and density of the mycelium. (C) Strain ESE00182 from the Cheese_2 population showing the fluffiness of the colony (D) First two PCA axes of the principal component analysis (PCA) of *G. candidum* strains based on their relative proportions of different volatile compounds produced. (E) Contribution of each volatile compound to the first two PCA axes. The compounds contributing the most to the differentiation were colored in red and labeled (i.e., those distant from 0 by an Euclidean distance >=0.1).

#### No adaptation to high salt concentration or milk origin in cheese populations

Cheese is a salty medium, with the percentage of salt varying from 0.5 g / 100 g for Emmental to 3 g / 100 g for Roquefort. Salt is added on the surface of cheeses to prevent the growth of contaminants, and cheese populations of *G. candidum* may thus have adapted to high salt concentrations. Cheeses display a wide range of salt concentrations so we tested four cheese media: unsalted, 1% salt as St Nectaire and cream cheeses, 2% as Camembert and goat cheeses and 4% as Roquefort blue cheeses. Wild populations grew faster than cheese populations in all salt concentrations tested, as on YPD and minimal media (Figure S10A ;Table S13).

Because all strains sampled from goat cheeses belonged to the Cheese_1 population, we tested whether this population was able to grow faster on goat cheese medium (1% salt) compared to other populations. We however found no significant interaction between population and media on radial growth effects, i.e. no specific adaptation to any particular kind of milk by the different populations (Figure S10B).

#### Contrasting volatile compound production between wild and cheese populations

Cheese ripening fungi, including *G. candidum*, contribute to cheese flavor through the production of volatile compounds ^56^. Flavor is a crucial criterion for cheese consumers and the cheese populations may have been selected for desirable and specific volatile compounds. We grew 14 *G. candidum* strains on a sterilized Camembert curd for 21 days at 10°C, *i.e.,* the ripening conditions of a Camembert. On average across compounds, the wild population produced five times higher quantities of volatiles than cheese populations. In order to compare the relative proportions of the different compounds, which is also an important aspect for flavor, we standardized the values by dividing all compound quantities by the total quantity of volatiles per sample. The PCA indicated a differentiation between wild and cheese strains in terms of volatile relative proportions (Figure 4D). The wild population thus produced combinations of volatile compounds different from cheese populations, with a high proportion of ethyl esters and ethyl acetates (Figure S10C). However, the impact of ethyl acetate on flavor is rather negative because it brings solvent type notes. In cheese, these esters are never predominant ^68,69^. Ethyl esters are involved in anaerobic metabolism and may be important for survival in the wild. By contrast, cheese strains produced many alcohols, ketones, aldehydes and sulfur compounds (Figure S10C), known for producing flavors typical of cheeses such as buttery, cheesy, fermented and aldehydic notes ^70^. These cheese-associated volatile compounds were present in similar absolute quantities in wild strains but were in minor proportions compared to other volatile compounds (Table S13), suggesting that cheese populations evolved a lower production of undesirable and unused volatiles. The overall balance between different volatile compounds is as important as volatile absolute quantities for flavor perception ^68^. The dimethyl sulfone, a compound previously reported being produced during the catabolism of L-methionine in *G. candidum*, is actually specifically produced by the cheese populations ^57,59^, with no difference between the cheese populations.

#### Cheese populations inhibit more the growth of food spoilers than wild populations

Cheese is a protein- and fat-rich medium, where many microorganisms, including desired microbes, but also spoilers, can thrive and thus compete for nutrients; for example, iron is limiting in cheese ^58,71,72^. Cheese *G. candidum* populations may have been selected for excluding competitors by inhibiting their growth ^50^. This fungus is known to inhibit fungal and bacterial food spoilers, such as *Aspergillus* species and *Listeria monocytogenes,* but these inhibitory activities have only been investigated in cheese *G. candidum* strains so far ^73–75^. We therefore tested whether cheese populations displayed better growth inhibition abilities than the wild population, using common fungal food spoilers as competitors: *Debaryomyces hansenii, Penicillium biforme, P. roqueforti* and *Scopulariopsis asperula*. We also tested whether growth inhibition of challengers occurred via secreted and/or volatile compounds. ***Inhibition by a mycelium lawn -*** In the first experiment, we grew challengers in a central spot for 24h, alone or after spreading out *G. candidum* to let it grow as a lawn; growth inhibition could occur in this setting by secreted molecules in the medium, volatile compounds and/or a physical barrier to reach nutrients. The growth of *D. hansenii* was completely inhibited by all populations of *G. candidum*. *Penicillium roqueforti* was strongly inhibited by *G. candidum*, in particular by the Cheese_2 population that completely prevented *P. roqueforti* growth (Figure 5A; Table S13). The growth of *Scopulariopsis asperula* and *P. biforme* was also inhibited by *G. candidum,* with a significant difference between competitor growth when spread alone or on a *G. candidum* lawn; the Cheese_2 population again inhibited better competitors than any other population (Figure 5A; Table S13).

**Figure 5:**
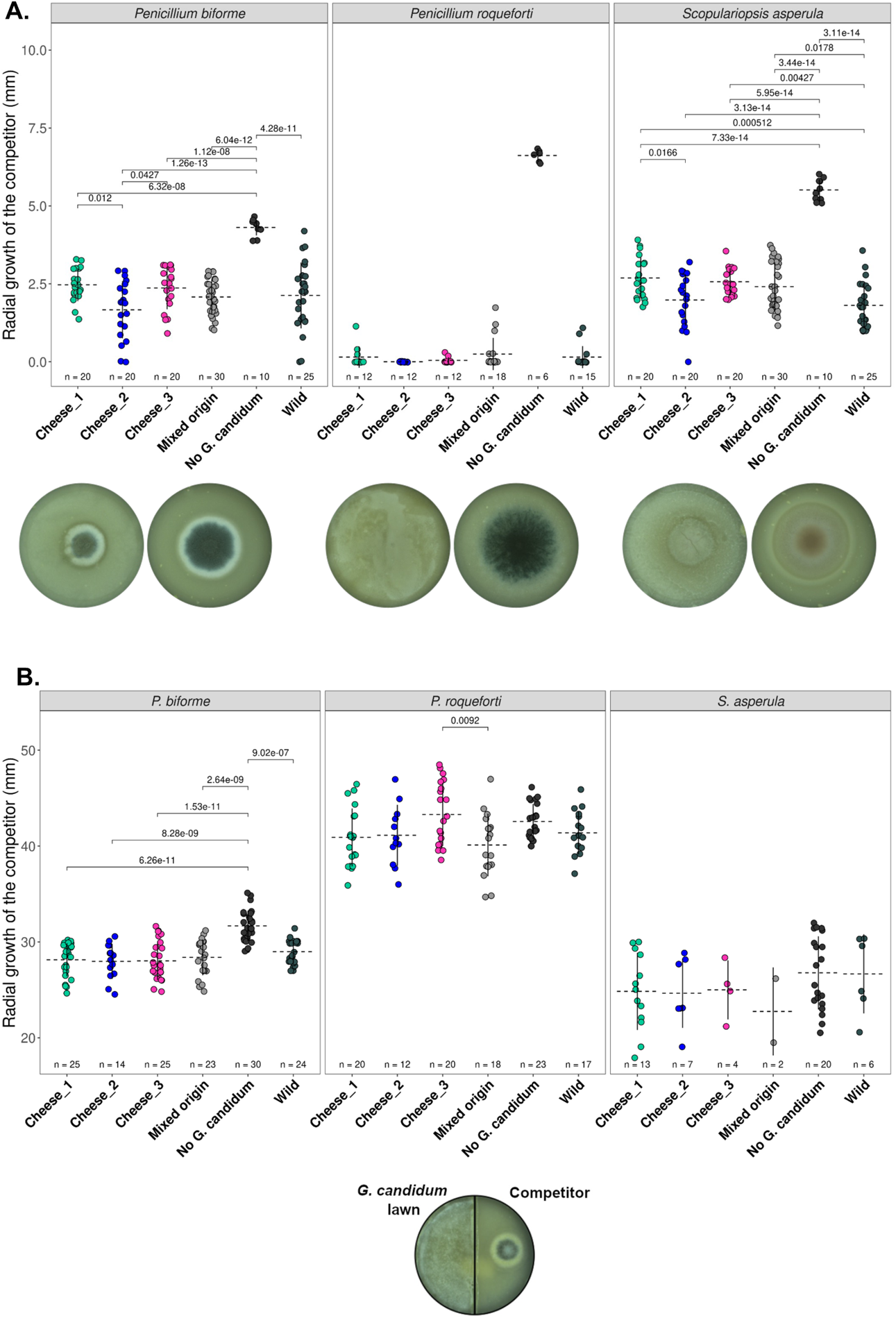
Competitive abilities of the different populations of *Geotrichum candidum* against *Penicillium biforme, P. roqueforti* and *Scopulariopsis asperula* challengers. (A) At the top, radial growth abilities of the competitors on lawns of *Geotrichum candidum* belonging to different populations (the three cheese populations, the mixed-origin and the wild populations). Each point represents a combination of the growth of a competitor strain on a lawn of a *G. candidum* strain. Horizontal dotted lines and vertical lines represent the mean and the standard deviation of the competitor growth in the population, respectively. The number *n* at the bottom of plots indicates the number of combinations of competitor-lawn used per population. The competitor was inoculated in a central point 24h later on a lawn of *G. candidum.* At the bottom, from left to right, are shown pictures of *P. biforme* ESE00023 on a *G. candidum* ESE00186 lawn and without any lawn, *P. roqueforti* ESE00645 on a *G. candidum* ESE00186 lawn and without any lawn, and *S. asperula* ESE01324 on a *G. candidum* ESE00198 lawn and without any lawn, all on a salted cheese medium. B: At the top, radial growth abilities of competitors, with various *G. candidum* strains belonging to different populations being grown on the other side of splitted Petri dishes. The competitor was inoculated in a central spot on one side and the *G. candidum* strain was spread on the other side of the splitted Petri dish (a picture is shown as an example below the figure). The medium is not contiguous between the two sides of Petri dishes, so that inhibition can only occur by volatile compounds. Horizontal dotted lines and vertical lines represent the mean and the standard deviation of the competitor growth in the focal population, respectively. The number *n* at the bottom of plots indicates the number of combinations of competitor-lawn used per population.

***Inhibition by volatile compounds -*** In a second experiment, we used splitted Petri dishes (the two parts being separated by a plastic barrier) to test whether cheese populations inhibited competitors to a greater extent than the wild population when only volatile compounds can reach challengers. No significant growth difference was observed between the growth alone and at the side of *G. candidum* for neither *S. asperula* nor *P. roqueforti* (Figure 5B, Table S13). Only *P. biforme* showed a significant growth inhibition by *G. candidum* in this setup (Table S13); such a growth inhibition by *G. candidum* from an isolated Petri dish compartment indicates that volatile compounds produced by *G. candidum* are able to impair the growth of some competitors.

## Discussion

### Domestication of the cheese-making fungus *Geotrichum candidum,* with three varieties displaying contrasting phenotypes

We found three differentiated clades based on genomic analyses of 98 *G. candidum* strains isolated from different kinds of cheeses, dairy products (e.g., raw milk), other food substrates (e.g., sausage) and other environments (e.g., plants). One clade was specific to cheeses, one to wild environments and one had mixed origins (dairy and other environments). Although the dairy environment was over-represented in our sampling, the 10 wild strains available captured a substantial diversity both in terms of substrates (e.g. soil, flower and polyurethane) and of geographic origins (i.e., Thailand, UK, French Guiana, Brazil, Egypt, Senegal, South Africa, Belgium, Spain and Sweden).

Within the cheese clade, we revealed the existence of three distinct genetic clusters and several admixed strains. The cheese_1 population included all strains isolated from goat and soft cheeses, such as Sainte Maure de Touraine or crottin de chavignol. The other cheese populations corresponded to strains isolated from other types of cheeses, such as pressed uncooked cheeses (e.g. Reblochon) or cooked (e.g. Comté) cheeses. This suggests that cheese-making practices have led to genetic differentiation by divergent selection. We did not detect in the present study any specific trait in the Cheese_1 population compared to the Cheese_3 population, but our experiments did not capture all aspects of cheese-making; in particular, a feature of goat cheeses caused by *G. candidum* that could be investigated is their convoluted aspect. Alternatively, the clustering of soft goat cheese strains may be due to migration by strain sharing among cheese makers with similar products, as shown for example in bread yeasts ^76^.

The cheese populations each had twice as low genetic diversity as the wild population, suggesting bottlenecks due to human selection. We did not, however, find that a single clonal lineage was used for most of cheeses, as is the case in *P. roqueforti* and in *P. camemberti* ^33,35^, indicating the lack of such extreme bottlenecks in *G. candidum*. The overall diversity in the cheese clade with the three populations pooled was even higher than in the wild population, which may be due to the relatively low number of wild strains available. Alternatively, the high diversity in the cheese clade as a whole may be due to i) the diversification into three populations under human divergent selection, ii) regular migration from a wild pool thanks to the natural occurrence of *G. candidum* in raw milk, thus preserving the diversity of strains isolated from cheeses, or iii) the divergence of each of the three cheese clusters from distinct but unsampled wild populations. According to this third hypothesis, the three cheese clusters would correspond to a diversification event that predated domestication, with selection of strains for cheese making from three already differentiated populations. In fact, the cheese clade was not nested within the wild clade, as could be expected if we had sampled the population-of-origin of the cheese clade. The traits in the studied wild clade may therefore not correspond exactly to those of the ancestral population(s) of the cheese clade.

We nevertheless found some evidence of domestication in the cheese clusters. We detected genomic footprints of adaptation to cheese, with the presence of genomic islands of differentiation and selective sweeps in cheese populations, in particular on a gene involved in iron uptake, that is limiting in cheese ^72^, and on a metalloprotease involved in casein degradation ^56^. The cheese populations further lacked genes not required in the human-made environment, i.e. tandem beta lactamase-like genes. The cheese populations may therefore have lost these genes due to relaxed selection, although we cannot exclude a gain in the wild clade. The cheese populations also carried a higher load of transposable elements, which may result from an expansion following relaxed selection; we cannot rule out a reduction in repetitive elements in the wild clade instead, although it seems less likely.

We also found evidence of phenotypic adaptation to cheese making in *G. candidum* cheese populations, with shared traits specific to cheese populations, beneficial for cheese making and different from wild and mixed-origin populations. Cheese populations produced a greater amount of volatiles typical of cheese aromas and attractive to consumers compared to wild strains. The cheese populations also displayed slower growth on all media, even cheese, and lower proteolysis activity, which may prevent a too fast degradation of products during maturation, as found in the Roquefort *P. roqueforti* population ^35^ and in the dry-cured meat *Penicillium* fungi ^77^. Alternatively, the slower growth could represent degeneration due to the accumulation of deleterious mutations because of genetic drift. However, the genetic diversity is not that low in the cheese clade, and slower growth seems a feature of cheese fungi ^78^. These shared traits may have been acquired in their common (already domesticated) ancestor, or represent convergence in the case of repeated domestication from three different wild populations.

We did not detect higher salt tolerance or higher lipolysis ability in cheese populations. Higher salt tolerance and lipolysis rates have been reported in the non-Roquefort *P. roqueforti* population compared to non-cheese populations, but not in the Roquefort *P. roqueforti* population ^35,79^. The dry-cured meat *Penicillium* fungi even had lower lipolysis rates ^77^. This may be due to a lack of selection if fast growth of fungi is not beneficial for cheese making, for example if it leads to degraded products. There may also be evolutionary constraints for salt adaptation. Another hypothesis is that selection has not been efficient enough for changing multiple traits at the same time, and/or because of migration. As *G. candidum* is naturally present in raw milk, regular migration may occur between cheese and wild populations in artisanal cheeses that do not use commercial starters. We did not detect evidence of wild-to-cheese gene flow except in the admixed strains, but we may not have identified all wild populations.

Within the cheese clade, we found phenotypic differences between the three populations, in terms of colony density, volatile compounds produced and the ability of competitor inhibition. They thus correspond to different varieties in the sense of genetically differentiated clusters with contrasting phenotypes. These differences may result from selection for different traits, for example for making different cheese types or cheese-making practices. Alternatively, but not exclusively, part of the differences may correspond to pre-existing differentiated traits if the three cheese populations derived from different, unsampled, wild populations.

The *G. candidum* Cheese_2 genetic cluster was phenotypically the most differentiated, with in particular a denser and fluffier mycelium and a higher competitive ability against challengers compared to the other populations. The Cheese_2 population had a stronger inhibition ability than the other *G. candidum* populations when molecules could diffuse in the medium and the mycelium could act as a barrier. The Cheese_2 population was also the most fluffy population (Figure 6) and had a beta-glucanase gene under positive selection. Therefore, the challenger inhibition of Cheese_2 may be due to either mycelium density as a physical barrier or degradation of competitor cell walls.

**Figure 6:**
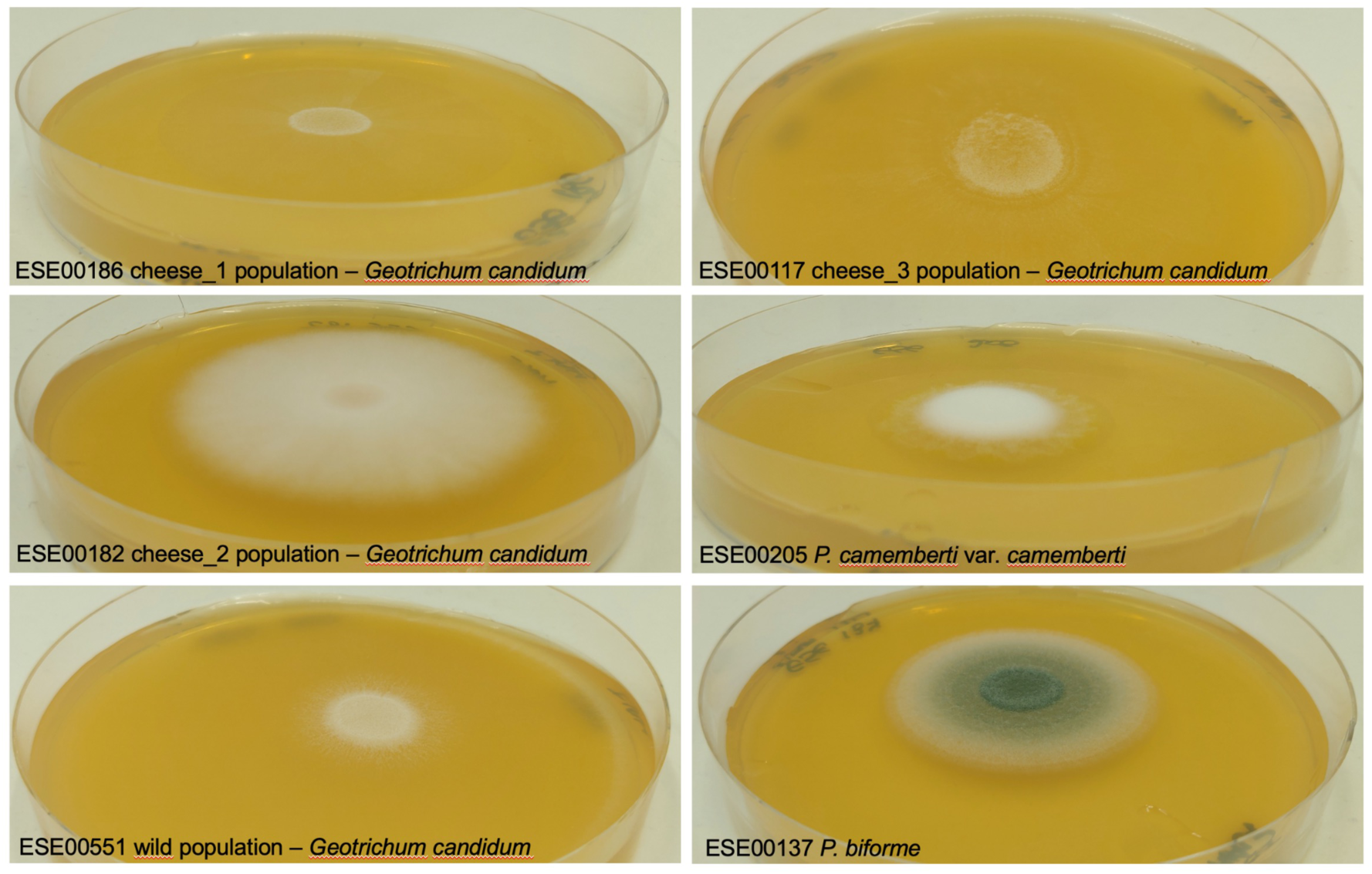
Illustration of colony phenotypes of the different *Geotrichum candidum* populations, *Penicillium camemberti* var. *camemberti* and *P. biforme*. A representative strain of each population of *Geotrichum candidum* as well as of *Penicillium camemberti* var. *camemberti* and *P. biforme* were grown on malt extract agar for seven days to illustrate the differences or convergence in mycelium color, density and fluffiness.

Our two sets of experiments enabled us to distinguish two mechanisms of competitor inhibition by *G. candidum,* and their action depended on the identity of the competitors. Indeed, *P. biforme* growth was inhibited by *G. candidum* in the splitted Petri dishes, suggesting that volatile compounds are able to impair its growth. On the contrary, *P. roqueforti* and *S. asperula* were only inhibited by *G. candidum* when grown together, i.e., when molecules could diffuse in their medium and *G. candidum* mycelium could form a physical barrier.

The genetic and phenotypic differentiation between the *G. candidum* clusters suggest that domestication occurred in several steps, with an ancient domestication event separating the mixed-origin and the wild clades, then the cheese and the mixed-origin clades, and yet more recently the three cheese clusters. Domestication also occurred stepwise in the *P. camemberti* clade ^33^: the first step has led to the domestication of *P. biforme* from the wild relative *P. fuscoglaucum,* and the second step to the divergence of *P. camemberti* from a *P. biforme*-like population. Such stepwise domestication has also been reported in other domesticated fungi, i.e. Cachaça yeasts, which are derived from wine yeasts ^18^ and also in crops ^12,80,81^. Alternatively, the divergence of the different cheese populations may have predated domestication; they would then correspond to unsampled wild populations, from which strains would have been independently isolated for cheese making.

Nevertheless, the Cheese_2 population appeared to represent a more advanced stage of domestication than the other cheese populations, with much lower genetic diversity, stronger genetic differentiation from other populations, and more differentiated traits from the wild population, with features beneficial for cheese making. The Cheese_2 population indeed likely corresponds to an asexually cultivated line for cheese making, displaying low diversity, a single mating type, a high level of linkage disequilibrium, an absence of reticulation in the neighbor-net network and a non-significant PHI test. The Cheese_2 population further display traits resembling other cheese fungi, with a denser and fluffier mycelium, a higher competitive ability and a complete lack of proteolysis activity. This may thus correspond to a stronger domestication syndrome, strikingly resembling the *P. camemberti* case. Indeed, *P. camemberti* is also a clonal lineage, recently derived from a cheese population with less extreme phenotypes, and having evolved drastic changes, with a white and fluffy mycelium^33^ (Figure 6).

In contrast to the situations in *P. roqueforti* and *P. camemberti* ^33,35^, commercial starter strains did not represent a single clonal lineage, being instead scattered throughout the cheese clade, some of them being admixed and/or belonged to small clonal groups (i.e. small terminal clusters in the tree with no branch length). These clonal lineages have each been isolated from different cheese types and different French regions, indicating asexual multiplication due to strain culture and commercialization, and migration between cheese makers ^76^. The existence of admixed commercial starter strains, belonging to clonal lineages, suggests that admixed lineages have been selected for beneficial traits for cheese making. Hybrids have been selected for their particular traits beneficial for food production in other domesticated fungi, such as cold tolerance in the hybrid yeast *Saccharomyces pastorianus* used for the production of lager beer ^66^.

### Phenotypic convergence but contrasted genetic diversity patterns among cheese fungi

We found evidence of convergence between *G. candidum* and other cheese-making fungi. As seen above, dense and white mycelial growth leading to a fluffy aspect at the expense of less rapid radial growth has strikingly evolved both in the Cheese_2 *G. candidum* and the *P. camemberti* var. *camemberti* lineages ^33^, thus representing a convergent phenotype between two distantly related cheese fungi ^78^. *Geotrichum candidum* is increasingly inoculated in milk in the place of *P. camemberti* for industrial soft cheese production, as it provides the white and fluffy desired aspect without the disadvantage of *P. camemberti* that browns the surface of Camembert cheeses at the end of the ripening process ^82^. Another convergence concerned proteolysis activity, which is low in *G. candidum* cheese populations as well as in the Roquefort *P. roqueforti* population, which may be crucial to avoid degraded cheeses ^35^. In terms of volatile compounds, we found specific and attractive flavors for cheese making in cheese strains, different from wild strains, as also documented in *P. roqueforti* ^35,38^.

In contrast to the striking convergence of phenotypes (Figure 4C), we found different patterns of diversity in *G. candidum* compared to *P. camemberti*. While *P. camemberti* and the non-Roquefort *P. roqueforti* populations are each a clonal lineage with virtually no genetic diversity ^33,35^, the *G. candidum* populations harboured each some diversity, even the less diverse, the Cheese_2 population. The Cheese_2 population of *G. candidum* indeed exhibited four times as low genetic diversity as the two other cheese populations, being of the same order of magnitude as in the Roquefort *P. roqueforti* population ^35^. The Cheese_1 population of *G. candidum* displayed the same level of diversity as the cheese-making species *P. biforme* and the Cheese_3 population as its wild relative *P. fuscoglaucum.* The different patterns of diversity may be due to different cheese-making practices, as *P. camemberti* and *P. roqueforti* are inoculated at the beginning of the cheesemaking process, while *P. biforme* and *G. candidum* are also naturally present in the cheese environment, either in raw milk or in barns.

The differentiation level between the whole cheese clade and the mixed-origin population in *G. candidum* was similar to that found between the domesticated *P. camemberti* mold and its wild closest relative species, *P. fuscoglaucum* (*FST* = 0.83; *dxy* = 6E-03), further supporting the view that the three cheese populations have been domesticated.

## Conclusion

Our study shows that fungi are excellent models to study domestication and independent adaptation events to similar environments and usage (Ropars and Giraud 2022). This is an important topic in evolutionary biology as it is important to understand whether independent adaptation events to similar environments leads to convergence in phenotypes, i.e., whether evolution is repeatable ^83–90^. We found here both similarities (convergence) and differences in the adaptation of *G. candidum* to cheese compared to other cheese fungi. One of the most striking convergent traits was the fluffy and white mycelium in *G. candidum* as in *P. camemberti* with a similar trade-off with radial growth ^33^. Greater competitive ability and lower proteolysis activity, as well as greater production of positive volatile compounds, also represent interesting convergent phenotypes between *G. candidum* and *Penicillium* cheese fungi, which are very distant fungal lineages. It will be interesting in future studies to investigate the genomic mechanisms underlying convergence, to assess whether the evolution of similar traits arose by similar or different *de novo* mutations, or by introgression or by the horizontal transfer of the same genomic regions, as already reported in *Penicillium* cheese fungi ^34,36^ and dry-cured meat *Penicillium* fungi ^77^.

Our findings also have industrial implications, revealing a high genetic diversity and genetic subdivision in a fungus widely used in the cheese industry, and the existence of genetically and phenotypically different populations used for cheese making, with specific and contrasted traits beneficial for cheese making. The most fluffy and most competitive cheese population corresponded to a clonal lineage which may represent the most recent and strongest selection event. The occurrence of recombination between cheese strains is highly relevant for cheese producers as it opens up possibilities for further improvement for cheese making. It is crucial to maintain a large genetic diversity in cheese *G. candidum* populations as this is essential for variety improvement and diversification and to avoid degeneration ^91^.

## Material and Methods

### Sampling

We isolated 53 strains from different kinds of cheeses (e.g. Camembert, Brie, Saint Nectaire, Ossau Iraty, comté, bleu de chèvre) from five European countries, Canada and the USA. Cheese crusts were left in the freezer for 24h to kill acarians. Then, we diluted a piece of each crust in sterile water and spread 50 uL of the suspension on a malt agar Petri dish. When colonies appeared on the Petri dish, typically after three days, we isolated the different morphotypes with a sterile toothpick and inoculated them on new Petri dishes. After seven days of growth, we performed monospore isolation by several dilution steps, in order to obtain separated colonies arising each from a single spore. We identified the species of these pure strains after DNA extraction by sequencing the 5’ end of the nuclear ribosomal large subunit (LSU rDNA) using the LROR/LR6 oligonucleotide primers ^92^. We also gathered 24 strains from INRAE, isolated from cheeses but also other environments (e.g. sand, hay, rainforest) and 15 strains from a French spore seller. We gathered all the wild strains available in public collections. For each strain, single spore cultures were generated to ensure the presence of a single genotype before DNA extraction.

The LMA-244 strain was inoculated on Yeast Extract Glucose (YEG) agar plates (10 g.L-1 of yeast extract (Fischer Scientific), 10 g.L-1 of D-glucose (EMD Chemicals) and 15 g.L-1 of Bacto agar (BD Diagnostics)) directly from 15% glycerol (v/v) stock cultures stored at -80°C. The plates were incubated in the dark for five days at 25°C.

#### DNA extraction, genome sequencing, assembly, annotation and mapping

We used the Nucleospin Soil Kit (Macherey-Nagel, Düren, Germany) to extract DNA from 88 *G. candidum* strains cultured for five days on malt agar. Sequencing was performed with Illumina HiSeq 3000 paired-end technology (Illumina Inc.), 2×150 bp. For the eight LMA strains, sequencing was performed using the Illumina HiSeq paired-end technology.

All Illumina reads were trimmed and adapters cleaned with Trimmomatic v0.36 ^93^. Leading or trailing low quality or N bases below a quality score of three were removed. For each read, only parts that had an average quality score higher than 20 on a four base window are kept. After these steps, only reads with a length of at least 36 bp were kept.

Cleaned Illumina reads were assembled with SPAdes v3.15.3 not using unpaired reads with “--careful” parameter.

For the LMA-244 strain, Genomic DNA was extracted using the Fungi/Yeast Genomic DNA Isolation Kit (Norgen Biotek Corp.) with the following modifications. Thirty milligrams of frozen grounded mycelium were thawed and homogenized in 500 µL of a 0.9% NaCl solution. The elution buffer was replaced by a Tris 10 mM buffer (pH 8). Following the extraction step, gDNA suspensions were purified and concentrated using Agencourt AMPure XP magnetic beads (Beckman-Coulter), according to the manufacturer’s protocol.

DNA concentration and purity were measured using a NanoDrop ND-1000 spectrophotometer (Thermo Fisher Scientific Inc., Wilmington, U.S.A.) and a Qubit Fluorometer 3.0 (Thermo Fisher Scientific Inc., Wilmington, U.S.A.).

The DNA library was prepared following the Pacific Biosciences 20 kb template preparation using BluePippin Size-Selection System protocol and the Pacific Biosciences Procedure & Checklist – Preparing Multiplexed Microbial Libraries Using SMRTbell Express Template Prep Kit 2.0 protocol. No DNA shearing was performed. The DNA damage repair, end repair and SMRT bell ligation steps were performed as described in the template preparation protocol with the SMRTbell Template Prep Kit 1.0 reagents and the SMRTbell Express Template Prep Kit 2.0 reagents (Pacific Biosciences, Menlo Park, CA, USA). The DNA library was size selected on a BluePippin system (Sage Science Inc., Beverly, MA, USA) using a cut-off range of 10 kb to 50 kb. The sequencing primer was annealed at a final concentration of 0.83 nM and the P6 v2 polymerase was bound at 0.50 nM while the sequencing primer was annealed with sequencing primer v4 at a final concentration of 1 nM and the Sequel 3.0 polymerase was bound at 0.5 nM.. The libraries were sequenced on a PacBio RS II instrument at a loading concentration (on-plate) of 160 pM using the MagBead OneCellPerWell loading protocol, DNA sequencing kit 4.0 v2, SMRT cells v3 and 4 hours movies.

Raw PacBio reads were corrected using Illumina reads already available and described in a previous article ^48^, with the default parameters of the LoRDEC software and trimmed with Canu v1.6 ^94,95^. Corrected and trimmed PacBio reads were then assembled using Canu v1.6. Illumina polishing of the Canu assembly was performed using Pilon v1.22 ^96^. A final assembly step was then performed with the hybrid assembler SPAdes v3.11.1 using the trimmed PacBio reads, the Illumina reads and the Pilon corrected assembly as trusted contigs ^97,98^. Additionally, the CLIB 918 assembly (Bioproject PRJEB5752) was used as a reference in the SPAdes script for the assembly of each *G. candidum* genome ^51^. Scaffolds were filtered using the khmer software with a length cut-off of 1,000 bp ^99^.

The LMA-244 PacBio assembly and reads have been deposited in GenBank: PRJNA482613. To annotate short read assemblies and the LMA-244 genome, gene prediction was performed using Augustus v3.4.0 ^100^. The training annotation file “saccharomyces” was used, with parameters as follows: “--gff3=on”, “--protein=on”, “--codingseq=on”, “--exonnames=on”, “--cds=on” and “--uniqueGeneId=true”. The output of Augustus and the CLIB 918 gff was provided to Funannotate v1.8.9 ^101^ for functional annotation. InterProscan was used under Funannotate pipeline locally ^102^. Funannotate then searched in the Pfam database v34.0 and dbCAN database version 10.0 with Hmmer v3.3.2 ^103–105^, in database UniProt version 2021_03 and database MEROPS version 12.0 with diamond blastp v2.0.11 ^106,107^, eggNOG-mapper v2 on the database eggNOG 5.0 ^108,109^.

Cleaned reads were mapped on the reference genomes CLIB 918 and LMA-244 using Bowtie2 v2.4.2 ^110^. Maximum fragment length was set to 1000 and the preset “very-sensitive-local” was used.

SAMtools v1.7 ^111^ was used to filter out duplicate reads and reads with a mapping quality score above ten for SNP calling and above one for CNV analyses.

In total, we have a dataset of 98 genomes, 88 being sequenced (Bioproject PRJNA866540), eight from the University of Laval (LMA strains: Bioproject PRJNA482576, PRJNA482605, PRJNA482610, PRJNA482613, PRJNA482616, PRJNA482619, PRJNA490507, PRJNA490528), one strain CLIB 918 from the Collection de Levures d’Intérêt Biotechnologique (Bioproject PRJEB5752), and one of the strain Phaff72-186 from the 1000 Fungal Genomes project (Bioproject PRJNA334358 NCBI).

### SNP calling

Single nucleotide polymorphisms (SNPs) were called using GATK v4.1.2.0 HaplotypeCaller, which provides one gVCF per strain (option -ERC GVCF). GVCFs were combined using GATK CombineGVCFs, genotypes with GATK GenotypeGVCFs, SNPs were selected using GATK SelectVariants (option -select-type SNP). SNPs were filtered using GATK VariantFiltration and options QUAL < 30, DP < 10, QD < 2.0, FS > 60.0, MQ < 40.0, SOR > 3.0, QRankSum < -12.5, ReadPosRankSum < -8.0. All processes from cleaning to variant calling were performed with Snakemake v5.3.0 (script available at https://github.com/BastienBennetot/Article_Geotrichum_2022).

### Phylogenetic analysis

We inferred phylogenetic relationships among the 98 isolates using the dataset of 699,755 SNPs in a maximum likelihood framework using IQ-Tree2 v2.1.1 ^112^. The tree has been midpoint rooted. The best-fit model chosen according to Bayesian information criterion (BIC) was TVMe+R2. Branch supports are ultrafast bootstrap support (1000 bootstrap replicates,^113^).

### Genetic structure

We used the dataset of 699,755 SNPs to infer population structure based on the mapping on the CLIB 918 reference genome. We used Splitstree v4.16.2 ^114^ for the neighbor-net analysis. We used the R package *Ade4* ^115–119^ for principal component analyses (PCA, centered and unscaled). We used NGSadmix v.33 ^120^ from the ANGSD ^121^ package (version 0.933-110-g6921bc6) to infer individual ancestry from genotype likelihoods based on realigned reads, by assuming a given number of populations. A Beagle file was first prepared from bam using ANGSD with the following parameters: “-uniqueOnly 1 -remove_bads 1 -only_proper_pairs 1-GL 1 -doMajorMinor 1 -doMaf 1 -doGlf 2 -SNP_pval 1e-6”. The Beagle file was used to run NGSadmix with 4 as the minimum number of informative individuals. Given the high number of strains genetically highly similar among cheese strains (that may represent clonal lineages), we randomly sampled one of the individuals for each group of clonemates identified on the ML tree as having fewer than 1,200 SNPs and filtered out the other strains (N=64 strains kept) to avoid biasing the analysis. The analysis was run for different *K* values, ranging from 2 to 10. A hundred independent runs were carried out for each number of clusters (*K*).

The nucleotide diversity *π* (Nei’s Pi; ^122,123^, the Tajima’s D, the Watterson’s *θ* ^124^, the fixation index *F_ST_* ^122^ and the absolute divergence *d_XY_* ^123^ were calculated using the *popgenome* package in R ^125^. Fixed, private and shared sites were counted using custom scripts available at https://github.com/BastienBennetot/fixed_shared_private_count, with bcftools version 1.11 (using htslib 1.13+ds). F3 tests were computed using the *admixr* package v0.9.1. The pairwise homology index (PHI) test was performed using PhiPack v1.1 and CLIB 918 genome as reference.

Linkage disequilibrium was calculated using vcftools v0.1.17 with the --hap-r2 parameter and a minimum distance between SNPs of 15,000 bp. Values were averaged when SNPs had the same distance.

Pairwise identity between an admixed strain and each non-admixed strain was calculated using overlapping sliding windows of 30 kb span and 5 kb step. Admixed clusters are indicated in Table S1. The custom script is available on https://github.com/BastienBennetot/Article_Geotrichum_2022

#### Copy number variation and identification of premature stop codons in CDS

Copy number variation (CNV) was analyzed using Control-FREEC v11.6 with the following parameters: ploidy was set to 1, non-overlapping windows of 500 bp, telomeric and centromeric regions were excluded, expected GC content was set between 0.25 and 0.55, minimum of consecutive windows to call a CNV set to 1. This analysis was performed using as references the CLIB 918 (cheese_3) and LMA-244 (wild) genome sequences. CNVs were classified in different groups when the median of copy number was different between populations. We defined three groups: regions for which copy number was different between wild and cheese populations, between mixed-origin and cheese populations and when at least one cheese population differed from another population. For each InterPro term present in these regions, we performed enrichment tests, i.e., a fisher exact test comparing the number of a particular InterPro term found in these regions and the whole genome (Table S9).

We used snpeff ^126^ to assess how each SNP affected the coding sequence of predicted proteins, in the vcf file containing all SNPs and all genomes of our dataset. We detected premature stop codons in the 7,150 CDS of the CLIB 918 genome and the 5,576 CDS of the LMA-244 genome using a custom script and bcftools v1.11.

### Analyzing the repeat landscape

In order to *de novo* detect repeats within *G. candidum*, RepeatModeler (v2.0.2 ^127^), using the ncbi engine (-engine ncbi) and the option -LTRStruct, was run on the pacbio genome assembly of LMA 244 generating a library of 176 repeats. The repeat redundancy was reduced using cd-hit-est, as described in Goubert et al., giving a final library of 108 repeats ^128^. To estimate the per strain copy number of each repeat, illumina reads were aligned using bwa mem (v0.7.17;^129^) to the repeat library and the median coverage for each repeat was then normalized by the LMA 244 genome wide median coverage.

### Detection of selective sweeps

We used SweeD ^130^ to detect selective sweeps based on site frequency spectra. We set the ploidy to 1 and the number of positions in the alignment where the CLR was computed to 1000 (gris option). We only kept the windows with the 1% highest likelihood per saffold for each cheese population.

### Detection of positive selection

The assemblies LMA-317, LMA-77 and LMA-563 have been excluded for this analysis because of a N50 under ten kb. All the 437441 predicted protein sequences from the 66 genomes of all cheese clades and mixed-origin clade were searched against each other with BLASTP using diamond v0.9.36 and clustered into orthologous groups using Orthagog v1.0.3 ^131^. For these analyses, we only kept single-copy orthologs shared between two populations. We compared the mixed origin population to each cheese population and the cheese clade. Multiple nucleotide sequence alignments with predicted gene sequences were then constructed using MACSE v2.0.3 with default parameters ^132^. We performed an approximative McDonald Kreitman tests using the R package PopGenome ^125^. The approximation comes from the fact that only codons with a single SNP are examined. The assumption of this version of the test is that the probability that two SNPs will appear in the same codon is very low. To identify genes evolving under positive selection in *G. candidum* genomes, ɑ, i.e. the representation of the proportion of substitutions driven by positive selection was used. Genes with an alpha under 0 were filtered out. Of these genes, only those with a Fisher’s test p-value under 0.05 were kept.

### Phenotypic characterization

#### Sampling and strain calibration

We used 36 *Geotrichum candidum* strains for laboratory experiments: seven from the Cheese_1 population, five from the Cheese_2 population, eleven from the Cheese_3 population, eight from the mixed-origin population and five from the wild population (Table S1). This set encompassed 26 strains isolated from dairies, one from other food environments and nine isolated from environments other than food. Experiments were initiated with spore suspensions calibrated to 1.10⁶ spores/mL with a hemocytometer.

#### Media preparation

All media were sterilized in an autoclave at 121°C for 20 minutes except those with cheese or milk for which the autoclave was run at 110°C for 15 minutes to avoid curdling. Each 94mm-diameter Petri dish was filled with 25mL of the appropriate medium. Cheese medium was prepared as follows for 800mL: 300g of unsalted cream cheese from La Doudou farm in Cheptainville, 16g agar, 8g NaCl dissolved in 200mL of deionized water. Deionized water was added to reach 800mL. pH was adjusted to 6.5 and drops of blue food dyes were added to enable fungal colony measures (white medium and white colonies are not distinguable). Yeast Peptone Dextrose (YPD) medium was prepared as follows for 1L: 10g Yeast extract, 10g Bacto Peptone, 10g glucose, 14g agar powder. Minimal medium was prepared as described in “Improved protocols for *Aspergillus* minimal medium: trace element and minimal medium salt stock solutions”, Terry W. Hill, Rhodes College, Etta Kafer, Simon Fraser University. Tributyrin agar was prepared as follows: Tributyrin medium 33 g/L, neutral Tributyrin 10 g/L, Bacto Agar 15 g/L. Ingredients were bought at Nutri-Bact company, Québec, Canada. Caseinate agar was prepared according to Frazier and Rupp, modified as follows: Calcium caseinate medium 37.2 g/L, Bacto Agar 15 g/L. Ingredients were bought at Nutri-Bact company, Québec, Canada. For yogurt media we used three different types of raw milk, i.e. sheep, goat and cow milks, coming from d’Armenon farm near Les Molières (Esonne, France), Noue farm in Celle les Bordes and Coubertin farm in Saint-Rémy-lès-Chevreuse respectively. Each medium was prepared following the same procedure: 1L of milk was mixed with 62.5g of Danone brand yogurt, heated for 5 hour at 43°C and stored in a fridge before use. A subset of 300g of this preparation was used with 16g of agar powder, 8g of NaCl, 4 drops of blue food dye and filled up with deionized water to reach 800mL.

#### Growth in different conditions and different media

Petri dishes were inoculated with 10µL of the 1.10^6^ cells/mL in a 10% glycerol solution. Inoculated Petri dishes were wrapped with plastic film before letting them grow in the dark. A millimeter rule was used to measure two opposite diameters of fungal colonies to estimate their growth. Means of these two measures were used for statistical analyses.

To test media and temperature effect on growth, *G. candidum* strains were grown on minimal, YPD and cheese media. We took pictures and measured their growth at seven, 11 and 14 days for minimal, YPD and cheese media at 10°C (ripening cellar temperature), at seven and 11 days for the cheese medium at 15°C and at seven days for minimal, YPD and cheese media at 25°C (Figure S8).

To test salt tolerance, *G. candidum* strains were grown at 10°C on cheese media of different salt concentrations: unsalted media, 1% salt as St Nectaire and cream cheeses, 2% as Camembert and goat cheeses and 4% as Roquefort blue cheeses. We took pictures and measured colony diameters after 14 days of growth.

To test adaptation of *G. candidum* populations to different milk origins, growth was measured on different yogurt media made from goat, sheep and cow raw milk for seven days at 25°C. To test lipolytic and proteolytic activities of *G. candidum* populations, we grew strains on tributyrin agar and caseinate agar, respectively. Each strain was inoculated in triplicate Petri dishes that were let grown at 25°C for 14 days. The radius of lysis was measured and the mean between triplicates was used for the analysis.

Pictures were taken using a Scan 1200 (Interscience). Petri dishes grown on cheese were analyzed using IRIS ^133^ which measured Integral opacity scores, defined as the sum of the brightness values for all the pixels within the colony bounds.

#### Volatile compounds analysis using Gas-chromatography mass-spectrometry (GC-MS)

Volatile compounds produced by *G. candidum* were analyzed using gas-chromatography mass-spectrometry (GC-MS). Compounds were extracted and concentrated by using a dynamic headspace (DHS) combined with a thermal desorption unit (TDU). Strains were grown for 21 days at 10°C (minimum Camembert ripening time) on a cheese agar medium made with Camembert-type curds. After 21 days, each Petri dish content, with its medium and *G. candidum* mycelium, was mixed with a fork for one minute, gathered in vials and immediately frozen in liquid nitrogen. For each sample, three grams of frozen cultured media were weighted and stored in vials with septum caps at -80°C. Sixteen hours before analysis, samples were stored at 4°C. The Cheese_2 population was not tested in this experiment because population delineation was not known at this time.

Dynamic headspace (DHS) conditions were as follows: Inert gas: He; Incubation: 30°C for 3min; Needle temperature: 120°C; Trap: nature tenax, 30°C, 450 mL He; He flow: 30 mL/min; Dry purge : temperature 30°C, 850 mL He, He flow 50 mL/ min. Thermal Desorption Unit (TDU) conditions were as follows: inert gas : He; Initial temperature: 30°C, then 60°C/min until 290°C kept for 7 minutes; Transfer temperature: 300°C. Cool Injection System (CIS) conditions were as follows: inert gas : He; Initial temperature: -100°C, then 12°C/s until 270°C kept for 5 minutes. Gas chromatograph (brand Agilent 7890B) was used with a polyethylene glycol (PEG) type polar phase column (HP-Innowax, ref. Agilent 19091N-116I, 60mx0.32mm, 0.25µm film thickness). Helium flow was set at 1.6mL/min. Samples were injected in splitless mode with a holding time of 1 minute. To optimize separation of compounds, a specific program of the gas chromatography oven was used, with initial temperature at 40°C for 5 minutes, rising temperature from 40°C to 155°C with a slope of 4°C/min, rising temperature from 155°C to 250°C with a slope of 20°C/min and then temperature was kept at 250°C for 5 minutes. A single quadrupole mass spectrometer was used to determine m/z of sample molecules (Agilent, référence 5977B MSD). Molecules were identified using NIST libraries (NIST 2017 Mass Spectral Library).

#### Competition experiments

To test the abilities of *G. candidum* populations to exclude other fungi by secreting molecules or volatile compounds, we compared the growth of competitors when grown alone and on a lawn of an already grown *G. candidum* mycelium. We inoculated a cheese medium with 150µL of a *G. candidum* calibrated spore solution (1.10⁶ spores/mL), spread evenly on the Petri dish. After 24h of growth, we inoculated 10µL of a competitor spore solution (1.10⁶ spores/mL) in a single spot, in the middle of the Petri dish. We used as competitors the following species and strains: *Penicillium biforme* (ESE00018, ESE00023, ESE00125, ESE00222)*, Penicillium roqueforti* (ESE00645, ESE00925, LCP06040), *Scopulariopsis asperula* (ESE00044, ESE00102, ESE00835, ESE01287, ESE01324) and *Debaryomyces hansenii* (ESE00284, ESE00561, ESE00576; Table S15). For each competitor, two Petri dishes were inoculated without any *G. candidum* as controls for measuring growth without a lawn.

We took pictures of the Petri dishes at 6 days, when the competitor mycelium grown alone was near the Petri dishes border; we measured colony size at 7 days for *P. biforme* and *P. roqueforti* and at 19 days for *D. hansenii*, which grows more slowly.

To test the abilities of *G. candidum* populations to exclude other microorganisms by producing volatile compounds, we set up an experiment with splitted Petri dishes where only air can be shared between the two parts. In one part of the Petri dish, we spread 75µL of a *G. candidum* spore solution (1.10⁶ spores/mL) and let it grow during 24 hours before adding on the other part of the Petri dish a drop of 5µL of a competitor spore solution (1.10⁶ spores/mL). For each competitor, two Petri dishes were inoculated without any *G. candidum* as controls. We used as competitors the following species and strains: *Penicillium biforme* (ESE00018, ESE00023, ESE00125, ESE00222, ESE00423)*, Penicillium roqueforti* (ESE00250, ESE00631, ESE00640, ESE00925) and *Scopulariopsis asperula*(ESE00044, ESE00102, ESE00835, ESE01287, ESE01324; Table S15). Petri dishes were grown at 10°C, measured and pictured at 11 days for *P. biforme* and *P. roqueforti* and 19 days for *Scopulariopsis asperula*.

#### Graphics and statistical analyses

Plots and statistical analyses were made using *ggplot2* ^134^, *rstatix* and *ggpubr* packages in the R environment. For ANOVAs, we used standard linear models in which all explanatory variables were discrete, with explained variables being radial growth for growth conditions (for media, temperature, salt content and adaptation to milk experiments), integral opacity score (for opacity experiment), relative proportions of volatiles compounds (for volatile compounds experiment) and radial growth of the competitor (for competition experiments). The explanatory variable common for all analyses was the ‘population’ of *G. candidum*. The variables ‘medium’, ‘day’ and ‘temperature’ were explanatory variables specific to the growth analysis. The ‘competitor species’ variable was specific to competition analyses. All variables and all interactions between them were implemented in the ANOVA and non-significant interactions were subsequently removed before performing post-ANOVA Tukey’s honest significant difference (HSD) tests. The data normality of residuals was checked; when residues deviated from normality (only for the opacity experiment), we also ran non-parametric tests (Wilcoxon ranking tests) using R. Radius of lysis for lipolytic and proteolytic activities experiments was often discrete, strains either showing lytic activity or not at all. This is why we decided to transform these data into qualitative discrete data in order to fit a generalized linear model with a binomial function as logit. Growth time (7, 14 and 21 days) and temperature (15 and 25°C) were taken as random variables because no fit could be achieved with little data and we wanted to test for population effect. Tukey contrasts were used to compare population means of populations when population effect was significant.

## Supporting information

Main figure

## Acknowledgments

We thank everyone who sent cheese crusts and Riwanon Lemee for providing 15 strains from Laboratoires STANDA, Caen, France. We thank Laura Prugneau, Jérémy Raynaud, Thomas Mari, Véronique Bougie and Jules Larouche for their help in the laboratory experiments. This work was funded by the Artifice ANR-19-CE20-0006-01 ANR grant to J.R. In addition, a NSERC-Discovery Grant supported the scholarship of V.P and data generation, and is held by S. Labrie (RGPIN-2017-06388).

## Author contributions

J.R. designed and supervised the study, and obtained funding. B.B., V.P., J.R. and A.S. generated the data. C.G. provided strains from the CIRM-Levures INRAE collection. B.B., J.-P.V., R.C.d.l.V. and S.O. analyzed the genomes. B.B., S.H., J.R. and A.S. performed the experiments. B.B. analyzed the data from laboratory experiments. M.H.L. and St.L. supervised the lipolysis and proteolysis analyses. B.B., So.L. and A.-C.P. performed the volatile compound experiment. T.G. contributed to interpretation; B.B., T.G. and J.R. wrote the manuscript, with contributions from all the authors.

## Supplementary Figure legends

**Figure S1: Population structure of *Geotrichum candidum*.**

Population subdivision inferred for *K* populations ranging from two to six. Colored bars represent the coefficients of membership in the K gene pools based on genomic data. Each bar represents a strain, its name being indicated at the bottom of the figure. The new color for each *K* increment is indicated on the right part. The second order rate of change in the likelihood (ΔK) peaked at *K*=6.

The names of the five populations considered in experiments are indicated on the last rows. Below the admixture plot, two rows indicate the type of milk from which the strains were sampled and the population subdivision.

**Figure S2: Pairwise identity between admixed and other strains, averaged per population of *Geotrichum candidum*.**

Pairwise identity along the genome, averaged per population, with only the first scaffold of the CLIB 918 genome shown

**Figure S3: Linkage disequilibrium against distance between SNPs for the five *Geotrichum candidum* populations.**

The r² is shown, representing the square of the correlation coefficient between two indicator variables; r² varies between 0 when two markers are associated at random and 1 when they provide identical information.

**Figure S4: Single nucleotide polymorphism (SNPs) inducing a premature stop codon for each *Geotrichum candidum* strain.**

We only show Stop -inducing SNPs present in more than three strains. Columns represent strains, ordered according to the maximum likelihood (ML) tree. Cells are colored in black when the corresponding SNP induces a premature stop codon, white when there is no substitution for this site compared to the reference genome and grey when the SNP status could not be assessed, substitution that induces other effects on the protein sequence were not present in this subset of Stop-inducing sites. Each row is a site; when multiple SNPs induced stop codons in the same gene, the corresponding rows were grouped and separated from other genes by black lines. The analysis was done using either the CLIB 918 (Cheese_3) genome (A) or the LMA-244 (Wild) genome (B) as a reference.

**Figure S5: Density of transposable element copy number relative to the LMA-244 strain.**

To better show the fat tail distribution, the y-axis (density) was cut at 25%. A red dashed line indicates the threshold of five times more copy number as the LMA-244 strain.

**Figure S6: Distributions of absolute divergence (dxy) values for different pairs of populations.**

Distribution of absolute divergence (*d_xy_*) values for each pairwise population comparison from the genomic scan analysis. Density is given as an overlapping window number for a specific value of *d_xy_*, each window being 7.5 kb wide with a step of 5 kb (optimal values based on variants densities). A black vertical line indicates the threshold of 1% highest values kept for the enrichment test.

**Figure S7: Genomic scan of within-population genetic diversity and between-population differentiation in *Geotrichum candidum*.**

Genomic of the nucleotide diversity *π*, watterson’s theta *θ_w_*, absolute divergence *d_xy_* and fixation index *F_ST_* ^122^ along the scaffold 1 of the CLIB 918 reference genome. At the bottom, a guide indicates genic regions in black and non genic regions in white. On the bottom of the figure, genic regions are shown in black (not positively selected) or red (positively selected in the McDonald and Kreitman test) rectangles. On the first panel (nucleotide diversity π), 5% lowest π values in the three cheese populations were highlighted by black dots. On the third panel (absolute divergence *d_xy_*), the 1% highest values of *d_xy_* of each pairwise comparison are highlighted by back dots.

**Figure S8: Differences in growth between the five populations of Geotrichum candidum populations for different media and temperatures.**

Mean radial growth of the three cheese populations, mixed-origin and wild populations on cheese (1% salt), yeast peptone dextrose (YPD) and minimal media at 10, 15 and 25°C for 7, 11 and 14 days.

Each point represents a strain, horizontal dotted lines and vertical lines represent the mean and the standard deviation of the population, respectively. The number *n* at the bottom of plots indicates the number of strains per population used for measuring these phenotypes. Significant pairwise Tukey tests are indicated with brackets and p-values.

**Figure S9: Differences in lipolytic and proteolytic activity among the five populations of Geotrichum candidum populations for different growing time and temperature**

A: Lipolytic activity of *G. candidum* at 15 and 25°C and grown for 7, 14 and 21 days. B: Proteolytic activity of *G. candidum* at 15 and 25°C and grown for 7, 14 and 21 days.

Each point represents a strain, horizontal dotted lines and vertical lines represent the mean and the standard deviation of the population respectively. The number *n* at the bottom of plots indicates the number of strains per population used for measuring these phenotypes. Significant pairwise Tukey tests are indicated with brackets and p-values.

**Figure S10: Differences in salt tolerance, growth on different milk types and volatile compounds between the five populations of *Geotrichum candidum*.**

Each point represents a strain, horizontal dotted lines and vertical lines represent the mean and the standard deviation of the phenotype in the population, respectively. The number *n* at the bottom of plots indicates the number of strains used per population for measuring the corresponding phenotypes. The pairwise Tukey tests performed to assess whether there were mean differences between populations are indicated with brackets and their p-values are given.

(A) Mean radial growth at 10°C of the three cheese, mixed-origin and wild populations of *G. candidum* on cheese agar medium with different salt concentrations: unsalted, 1% salt for mimicking St Nectaire and cream cheeses, 2% salt for mimicking Camembert and goat cheeses and 4% salt for mimicking blue cheeses.

(B) Mean radial growth at 10°C of the three cheese populations, the mixed-origin and the wild populations on yogurt agar media made with raw cow, goat and sheep milks.

(C) Relative proportions of major volatile compounds in Cheese_1, Cheese_3, mixed-origin and wild populations of *G. candidum.* The volatile compounds shown were those contributing the most to the two first PCA axes or those that are important for cheese ripening. For each compound, the related corresponding descriptor from <u>thegoodscentscompany.com</u> was added.

## Supplementary Table legends

**Table S1: Description of the origin, population assignation, type of cheeses, sequencing statistics, and tested phenotypes of the Geotrichum candidum strains used in this study.**

Strains: Names of the strains; ESE (Ecology Systematics and Evolution) collection ID : collection ID of strains kept at the ESE laboratory; Population: population to which the strain was assigned in this study; Clonal group: strains with the same number are clonemates (fewer than 1,200 SNPs between them); Species name: species ID identified either based on public collection or from ITS identification for strains collected in this study; Type of cheeses: if the strain was isolated from a cheese, it indicates what kind of cheeses, i.e. soft natural rind, soft, pressed uncooked, pressed cooked or blue cheeses; Environment of sampling simplified: broader categories for source of sampling; Milk type: if the strain was isolated from dairy products, it indicates the animal species from which milk originated; Location: Region or country of origin; Mating type: mating type identified in *G. candidum,* either MATA or MATB^51^.

**Table S2: Population genetics statistics estimating genetic differentiation (F_ST_, d_XY_) between the five Geotrichum candidum populations and between populations in other cheese fungi (Penicillium camemberti and Penicillium roqueforti)** ^33,35^

Cells are colored from the lowest (white) and the highest (red) value of each indices.

**Table S3: Proportions of fixed, shared and private SNPs for each pair of populations of Geotrichum candidum population and of two other cheese fungi (Penicillium camemberti and Penicillium roqueforti)** ^33,35^.

The method of attributions of SNPs to different categories are available on https://github.com/BastienBennetot/fixed_shared_private_count.

**Table S4: F3 test performed on each trio of populations of Geotrichum candidum populations.**

The F3 test is a test between three populations. It tests whether a target population (C) is admixed between two source populations (A and B) and gives a measure of shared drift between two test populations (A and B) from an outgroup (C). In case of introgression, we expect negative F3 values. A Z-score is computed based on F3 value and the standard error to assess the deviation from zero of the F3 value. If the Z-score is inferior to minus three then we can conclude a significant rejection of the Null hypothesis (F3 value is not negative).

**Table S5: Population genetics statistics estimating genetic diversity (𝜋, watterson’s θ and Tajima’s D) in the five Geotrichum candidum populations and populations of three other fungal species (Penicillium roqueforti, P. camemberti and Saccharomyces cerevisiae)** ^29,33,35^.

Cells are colored from the lowest (white) and the highest (red) value of each indices.

**Table S6: Distribution of the two mating types in each population and test of the deviation from a 1:1 ratio.**

**Table S7: Phi test of each population of *Geotrichum candidum* using the first scaffold of CLIB 918 assembly.**

The pairwise homoplasy index (PHI) test assesses with permutations the null hypothesis of no recombination by looking at the genealogical association among adjacent sites.

**Table S8: Number and percentage of single nucleotide polymorphism (SNPs) classified by impact, functional class, effect and genomic regions for each Geotrichum candidum population.**

Based on the 7,150 coding sequences (CDS) of the CLIB 918 assembly, the effect of all variants was assessed. Each variant is categorized in different functional effects on the protein sequence. Results are averaged per population and shown either as the mean number or percentage of SNPs. Results were computed using snpeff ^126^.

(A) Total number of single nucleotide polymorphisms (SNPs) within each *G. candidum* population.

(B) Putative variant impact prediction. Different impacts categories are defined in Snpeff manual in ‘Input & output files’ section

(C) Protein sequence effect of SNPs. Variants can either not change amino acid sequence (silent), change the amino acid (missense) or induce stop codons (nonsense)

(D) Functional effect of SNPs defined in Snpeff manual in ‘Input & output files’ section

(E) Position of the SNPs compared to genes

**Table S9: Copy number variants differentiating Geotrichum candidum populations and test for enrichment in gene ontologies contained within these windows.**

(A) Number of windows subsetted in each clade comparison using the LMA 244 assembly as reference

(B) Number of windows subsetted in each clade comparison using the CLIB 918 assembly as reference

(C) Enrichment test on gene ontologies (GO) present within the subsets of windows in each clade comparison using the CLIB 918 assembly as reference

(D) Enrichment test on the gene ontologies (GO) present within the subsets of windows in each clade comparison using the LMA 244 assembly as reference

**Table S10: Repeat copy number for the different strains of Geotrichum candidum.**

In order to *de novo* detect repeats within *Geotrichum candidum*, RepeatModeler v2.0.2 ^127^ was run on the pacbio genome assembly of LMA 244 generating a library of 176 repeats. The repeat redundancy was reduced using cd-hit-est, giving a final library of 108 repeats (presented in column “Clustering of repeat family”) ^128^. The type and family of these repeats is indicated when it could be inferred (column type of repeat and repeat family). To estimate the per-strain copy number for each repeat, illumina reads were aligned using bwa mem (v0.7.17; Li, 2013) to the repeat library and the median coverage for each repeat was then normalized by the LMA 244 genome wide median coverage. Strain: name of the strains analysed; relative median coverage: Coverage of all genomic reads mapped to the repeated sequence normalized by genome-wide coverage before taking the median of all nucleotides of the repeated sequence (gives an idea of the copy number of the repeated element genome-wide); Copy number relative to LMA-244 strains: relative median coverage of the strain divided by the one of LMA-244 (wild) strain.

**Table S11: Gene functions that showed footprints of divergent selection and recent selective sweeps.**

Gene functions that were detected either by keeping 1% highest of absolute divergence (dXY) between cheese and wild strains, or 5% lowest nucleotide diversity (𝜋) in the cheese populations but not in the wild population, or being in the 1% highest likelihood of being under selective sweep (based on sweeD analysis) were compared for each cheese populations. Windows were 7.5 kb wide and overlapping windows with a step of 5 kb for the dXY and the 𝜋 analysis.

**Table S12: Results of the McDonald and Kreitman (MK) test for positive selection.**

Genes evolving under positive selection were assessed using McDonald and Kreitman (MK) tests. Using contrasting levels of polymorphism and divergence at neutral and functional sites, MK tests contrast the fraction of substitutions at the functional sites that were driven by positive selection. When the p value was lower than 0.05, we considered that the gene was under positive selection. Only genes under positive selection are shown in the Table. The mixed-origin population was compared to the cheese clade (A), Cheese_1 (B), Cheese_2 (C) and Cheese_3 (D), and wild population was compared to the mixed-origin and the cheese populations as a whole (E).

Ortho_id: identifier of the orthologous gene; P1_nonsyn: number of non-synonymous polymorphisms in the first population; P2_nonsyn: number of non-synonymous polymorphisms in the second population; P1_syn: number of synonymous polymorphisms in the first population; P2_syn: number of synonymous polymorphisms in the second population; D_nonsyn: number of non-synonymous substitutions; D_syn: number of synonymous substitutions; neutrality.index: quantifies the degree of departure from neutrality; alpha: proportion of substitutions driven by positive selection; fisher.P.value: P-value of the MK test; GeneID: gene identifier in the CLIB 918 assembly annotation; Contig Start: Start of the gene sequence; Stop: Stop of the gene sequence; Name: Name of the closest orthologous annotated genes; Product: Function of the protein; PFAM: pfam database annotation; InterPro: InterPro database annotation

**Table S13: Analysis of variance (ANOVA) of all phenotypes and post-hoc tests.**

All outputs of analyses of variance (ANOVA) (A) and post-hoc tests (B) based on linear models that were used for phenotypic analyses. Linear models tested: Effect of media, temperature on radial growth (1); effect of media and population on radial growth of *Geotrichum candidum* at 25°C (2); effect of salt content and population on radial growth (3); effect of milk origin and population on radial growth (4); effect of media and population on opacity (5); effect of population of *G. candidum* on competitor growth (6); competition abilities by volatiles on splitted Petri dishes (7); effect of population on relative proportions of volatile compounds (8).

In ANOVA table, columns correspond to degree of freedom (Df), sum of squares (Sum Sq), mean square (Mean Sq), the F statistic (value) and the p-value of this test.

Columns in post-hoc Tables before the “term column” gives the condition kept for each single test. term: variable used for pairwise comparison; group1: the group compared against group2; group2: the second group, compared to group1; null.value: value of group1 - group2 under the null hypothesis; estimate: value of group1 - group2 using the data; conf.low: Lowest value of the confidence interval; conf.high: highest value of the confidence interval; p.adj: adjusted p-value; p.ad.signif: significance of the adjusted p-value (p-value >0.05:n.s.; <0.05:*; <0.01:**; <0.001:***, <1E-04:****). Post hoc test for assessing the effect of salt content and population on radial growth of *G. candidum*.

**Table S14: Analysis of variance (ANOVA) and post-hoc tests of lipolysis and proteolysis experiments.**

All outputs of analyses of variance (ANOVA) (A) and post-hoc tests (B) based on the different linear models that were used for lipolysis (1) and proteolysis (2) analyses. Radius of lysis for lipolytic and proteolytic activities experiments was often discrete, strains either showing lytic activity or not at all. Thus, data were transformed into qualitative discrete data and a generalized linear model with a binomial function as logit was fitted. No post-hoc tests were computed for the lipolysis analysis because there was no population effect.

In the ANOVA table: σ2: mean random effect variance of the model; *τ00*: random intercept variance of a given variable, or between-subject variance; ICC: intraclass correlation coefficient; N variable: number of categories of the given variable; Observations: sample size of the model; marginal R^2^: variance explained by the fixed effects; conditional R^2^: interpreted as the variance explained by the entire model (i.e. the fixed and random effects).

In the post hoc table: Linear Hypotheses: null hypothesis considered for each post-hoc test; Estimate: Measured value; Std. Error: standard error of this measure; z value: Z statistic of this test.

**Table S15: Description of the origin and species of strains used in the competition experiments.**

ESE collection ID: Author collection ID for this strain; Previous name in public collections: name of the strain in other public collections; Species: Species of the strain; Origin of sampling: Environment of sampling for this strain; Milk (if cheese): species from which the milk originated for the dairy product from which the strain was sampled; Location of sampling; geographical location of the sampling; Cheese shop: where the cheese was bought for cheese strains; Information on sampling date: Sampling date when known.

## Notes

https://github.com/BastienBennetot/Article_Geotrichum_2022

### Competing Interest Statement

The authors have declared no competing interest.

### Summary of Updates

Adding the badge and a link to the PCI evolutionnary biology recommandation

