## Supplementary material for "Domestication of different varieties in the cheese-making fungus *Geotrichum candidum*": Main figure

Figure 1

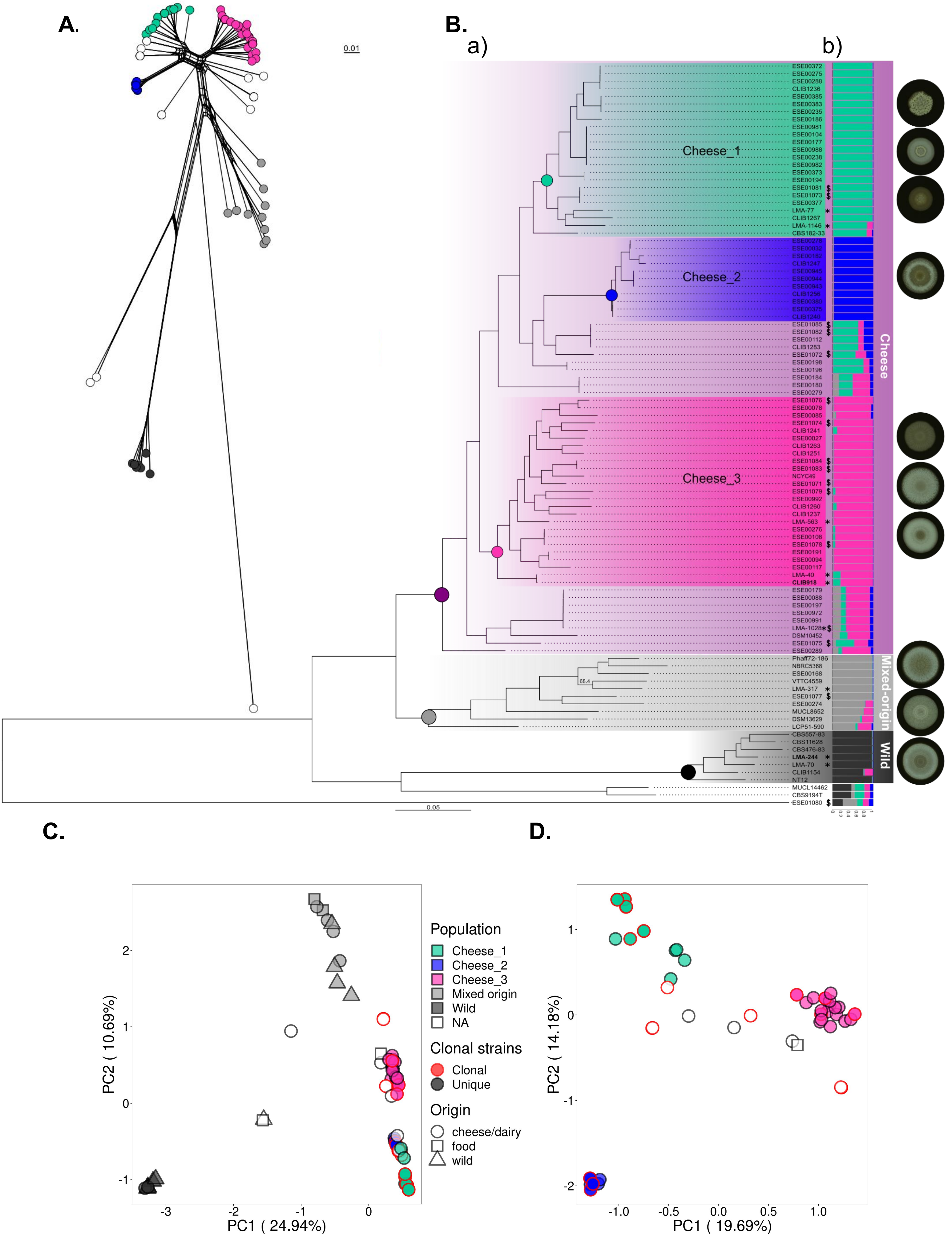

A.
0.01
B.
a)
b)
Cheese_1
Cheese_2
Cheese_3
Cheese
Mixed-origin
Wild
0.05
C.
D.
PC1 ( 24.94%)
PC2 ( 10.69%)
PC1 ( 19.69%)
PC2 ( 14.18%)
Population
Cheese_1
Cheese_2
Cheese_3
Mixed origin
Wild
NA
Clonal strains
Clonal
Unique
Origin
cheese/dairy
food
wild

Figure 1: Phylogenetic relationships and population structure of 98 strains of *Geotrichum candidum,* based on whole-genome data

(A) Neighbor-net analysis based on a single nucleotide polymorphism (SNP) distance matrix. The scale bar represents 0.01 substitutions per site for branch lengths.

(B) Genetic relationships among strains and population structure in *G. candidum* based on 699,755 SNPs. a) Maximum likelihood tree showing phylogenetic relationships among the 98 isolates used in this study. All nodes are supported by bootstrap support >98% (bootstrap analysis with 1000 resampled datasets). The scale bar represents 0.05 substitutions per site for branch lengths. We used the midpoint rooting method to root the tree. The "$" symbol pinpoints commercial starter strains and "*" the PacBio sequences. Genomes used as references are written in bold. b) Population subdivision inferred for *K* = 5. Colored bars represent the coefficients of membership in the five gene pools based on SNP data.

(C) Principal component analysis (PCA) based on 699,755 SNPs and 98 strains. Genetic clusters are represented by the same colors on all panels: light blue for Cheese_1, dark blue for Cheese_2, pink for Cheese_3, light grey for the mixed-origin population and dark grey for the wild population. The symbols represent the environment from which strains were sampled: circle for cheese/dairy, square for food and triangle for wild environment. Symbols are circled in red when multiple strains are overlapping due to clonal lineages (with a threshold set at <1,200 SNP for defining clonal strains).

(D) PCA based on the 323,385 SNPs when the dataset was restricted to the 78 strains from the cheese clade.

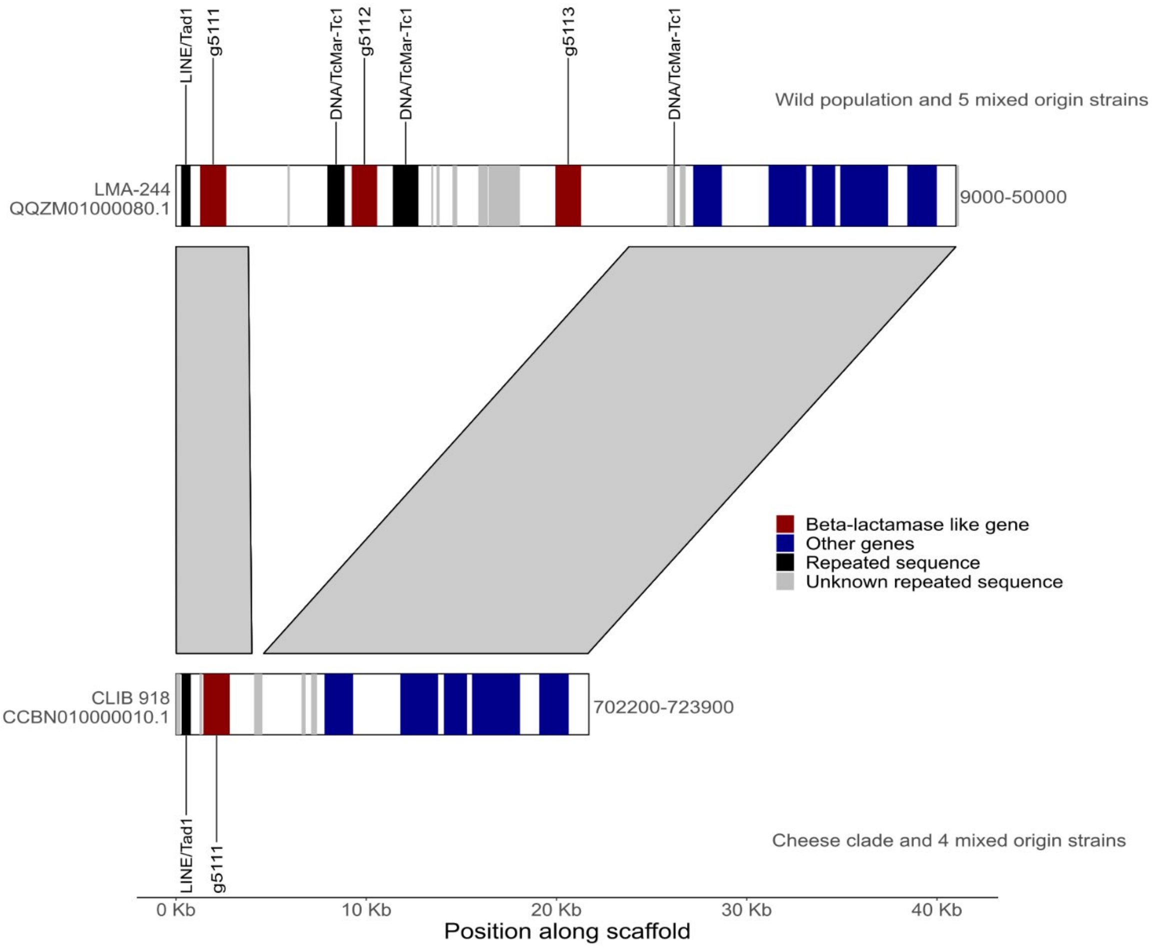


Figure 2: Lack of the beta lactamase-like genes in the cheese clade of *Geotrichum candidum*

Synteny between parts of the scaffold QQZM01000080.1 of the LMA-244 (wild) strain against the scaffold CCBN010000010.1 of the cheese CLIB 918 (Cheese_3) strain. Beta-lactamase-like genes are annotated in red while other genes are displayed in blue. Black triangles indicate positions with repeated sequences. All strains from the cheese clade and five strains from the mixed-origin populations (LMA-317, ESE00274, MUCL8652 and ESE00540) lacked the g5112 and g5113 genes, both encoding for beta-lactamase like.

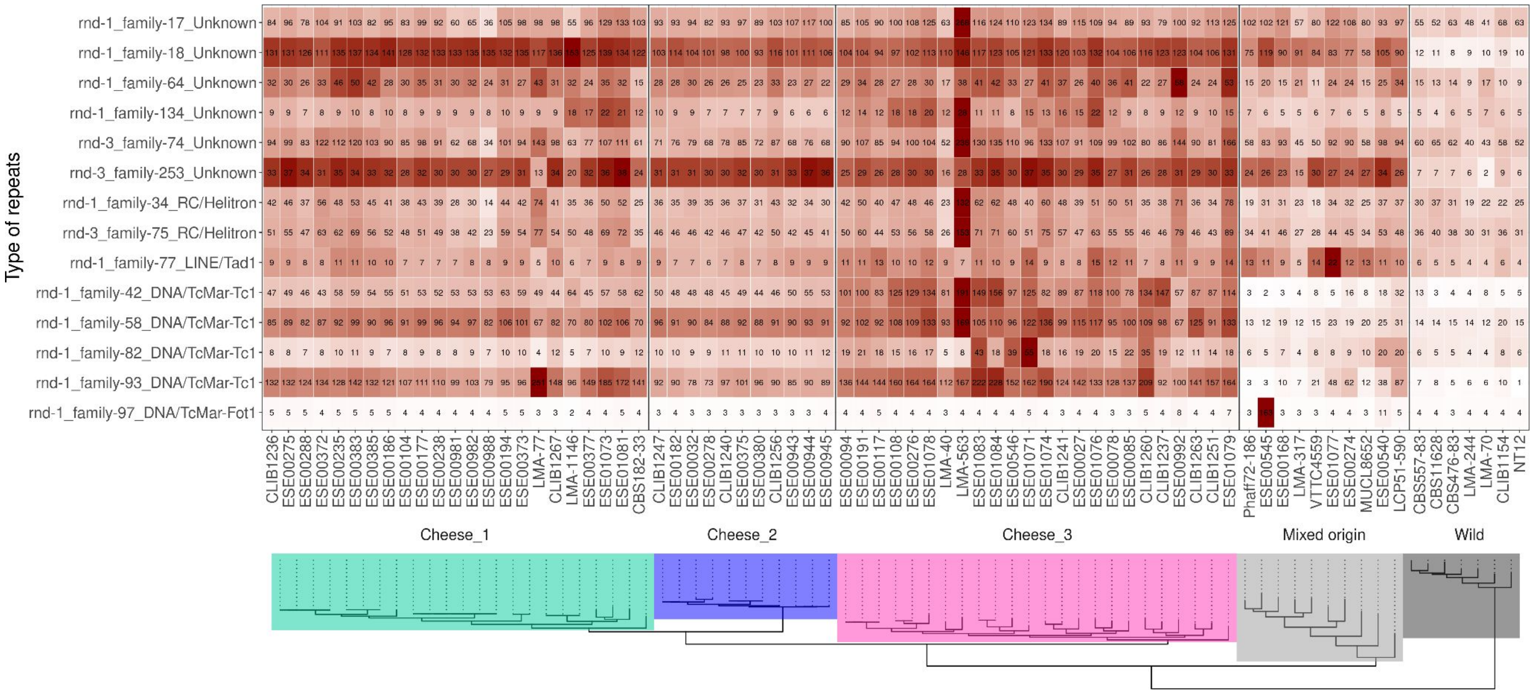


Figure 3: Heatmap of repeats in the cheese populations

The total number of copies is indicated in the center of each cell and by the grey to red color scale. The maximum likelihood (ML) tree from the figure 1 (without admixed strains) is plotted below strain names.

Figure 4

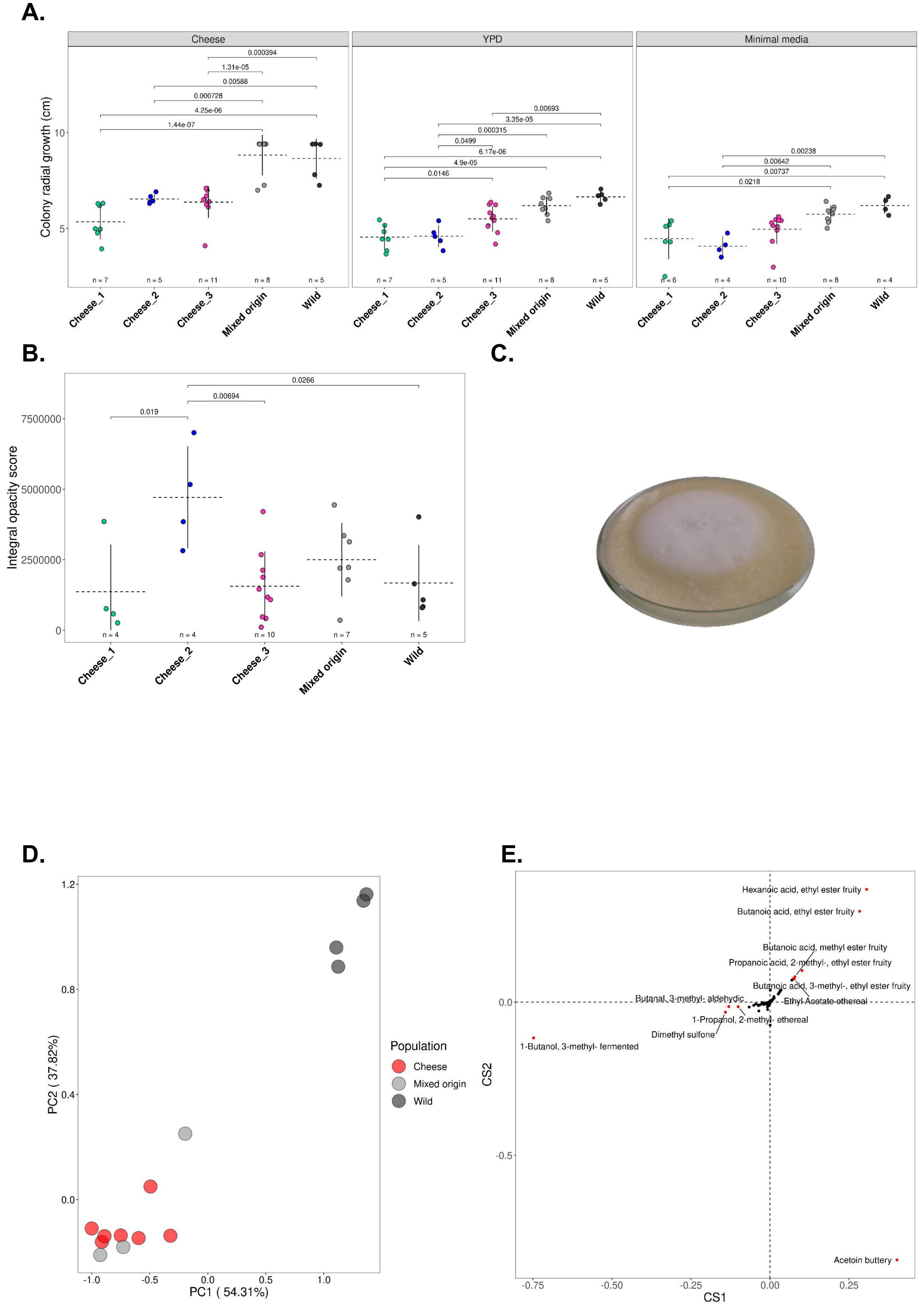
A.
Cheese
YPD
Minimal media
Colony radial growth (cm)
0.000394
1.31e-05
0.00588
0.000728
4.25e-06
1.44e-07
0.00693
3.35e-05
0.000315
0.0499
6.17e-06
4.9e-05
0.0146
0.00238
0.00642
0.00737
0.0218
n = 7
n = 5
n = 11
n = 8
n = 5
n = 6
n = 4
n = 10
n = 8
n = 4
Cheese_1
Cheese_2
Cheese_3
Mixed origin
Wild
B.
C.
Integral opacity score
0.0266
0.00694
0.019
7500000
5000000
2500000
0
n = 4
n = 4
n = 10
n = 7
n = 5
Cheese_1
Cheese_2
Cheese_3
Mixed origin
Wild
D.
E.
PC1 ( 54.31%)
PC2 ( 37.82%)
Population
Cheese
Mixed origin
Wild
Hexanoic acid, ethyl ester fruity
Butanoic acid, ethyl ester fruity
Butanoic acid, methyl ester fruity
Propanoic acid, 2-methyl-, ethyl ester fruity
Butanoic acid, 3-methyl-, ethyl ester fruity
Ethyl Acetate ethereal
Butanal, 3-methyl- aldehydic
1-Propanol, 2-methyl- ethereal
Dimethyl sulfone
1-Butanol, 3-methyl- fermented
Acetoin buttery
CS1
CS2

Figure 4: Differences in growth, opacity and volatile compounds between the five populations of *Geotrichum candidum*

Each point represents a strain, horizontal dotted lines and vertical lines represent the mean and the standard deviation of the phenotype in the population, respectively. The number *n* at the bottom of plots indicates the number of strains used per population for measuring the corresponding phenotypes. The pairwise Tukey tests performed to assess whether there were mean differences between populations are indicated between brackets with their p-values.

(A) Mean radial growth of the three cheeses, mixed-origin and wild populations on cheese (1% salt), yeast peptone dextrose (YPD) and minimal media at 25°C.

(B) Differences in opacity between the three cheese populations, mixed-origin and wild populations on cheese medium (1% salt) at 10°C. Integral opacity is defined as the sum of the brightness values for all the pixels within the fungal colony bounds and measures the whiteness and density of the mycelium.

(C) Strain ESE00182 from the Cheese_2 population showing the fluffiness of the colony

(D) First two PCA axes of the principal component analysis (PCA) of *G. candidum* strains based on their relative proportions of different volatile compounds produced.

(E) Contribution of each volatile compound to the first two PCA axes. The compounds contributing the most to the differentiation were colored in red and labeled (i.e., those distant from 0 by an Euclidean distance >=0.1).

Figure 5

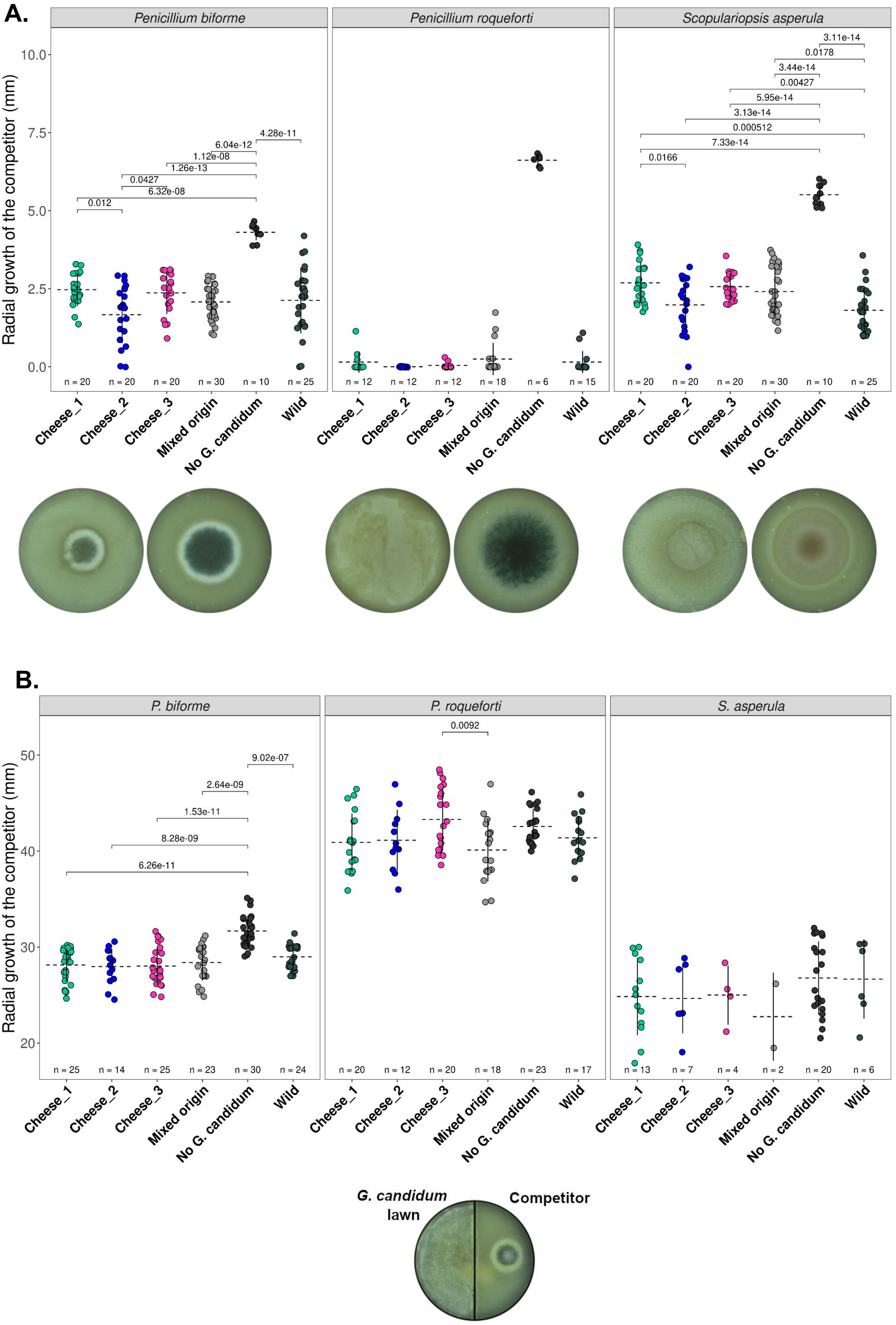

A.
Penicillium biforme
Penicillium roqueforti
Scopulariopsis asperula
Radial growth of the competitor (mm)
10.0
7.5
5.0
2.5
0.0
0.012
0.0427
6.32e-08
1.26e-13
1.12e-08
6.04e-12
4.28e-11
0.0166
7.33e-14
0.000512
3.13e-14
5.95e-14
0.00427
3.44e-14
0.0178
3.11e-14
n = 20
n = 20
n = 20
n = 30
n = 10
n = 25
n = 12
n = 12
n = 12
n = 18
n = 6
n = 15
n = 20
n = 20
n = 20
n = 30
n = 10
n = 25
Cheese_1
Cheese_2
Cheese_3
Mixed origin
No G. candidum
Wild
B.
P. biforme
P. roqueforti
S. asperula
Radial growth of the competitor (mm)
50
40
30
20
6.26e-11
8.28e-09
1.53e-11
2.64e-09
9.02e-07
0.0092
n = 25
n = 14
n = 25
n = 23
n = 30
n = 24
n = 20
n = 12
n = 20
n = 18
n = 23
n = 17
n = 13
n = 7
n = 4
n = 2
n = 20
n = 6
Cheese_1
Cheese_2
Cheese_3
Mixed origin
No G. candidum
Wild
G. candidum lawn
Competitor

Figure 5: Competitive abilities of the different populations of *Geotrichum candidum* against *Penicillium biforme, P. roqueforti* and *Scopulariopsis asperula* challengers.

(A) At the top, radial growth abilities of the competitors on lawns of *Geotrichum candidum* belonging to different populations (the three cheese populations, the mixed-origin and the wild populations). Each point represents a combination of the growth of a competitor strain on a lawn of a *G. candidum* strain. Horizontal dotted lines and vertical lines represent the mean and the standard deviation of the competitor growth in the population, respectively. The number *n* at the bottom of plots indicates the number of combinations of competitor-lawn used per population. The competitor was inoculated in a central point 24h later on a lawn of *G. candidum.* At the bottom, from left to right, are shown pictures of *P. biforme* ESE00023 on a *G. candidum* ESE00186 lawn and without any lawn, *P. roqueforti* ESE00645 on a *G. candidum* ESE00186 lawn and without any lawn, and *S. asperula* ESE01324 on a *G. candidum* ESE00198 lawn and without any lawn, all on a salted cheese medium.

B: At the top, radial growth abilities of competitors, with various *G. candidum* strains belonging to different populations being grown on the other side of splitted Petri dishes. The competitor was inoculated in a central spot on one side and the *G. candidum* strain was spread on the other side of the splitted Petri dish (a picture is shown as an example below the figure). The medium is not contiguous between the two sides of Petri dishes, so that inhibition can only occur by volatile compounds. Horizontal dotted lines and vertical lines represent the mean and the standard deviation of the competitor growth in the focal population, respectively. The number *n* at the bottom of plots indicates the number of combinations of competitor-lawn used per population.
