## Supplementary figure for "Domestication of different varieties in the cheese-making fungus *Geotrichum candidum*"

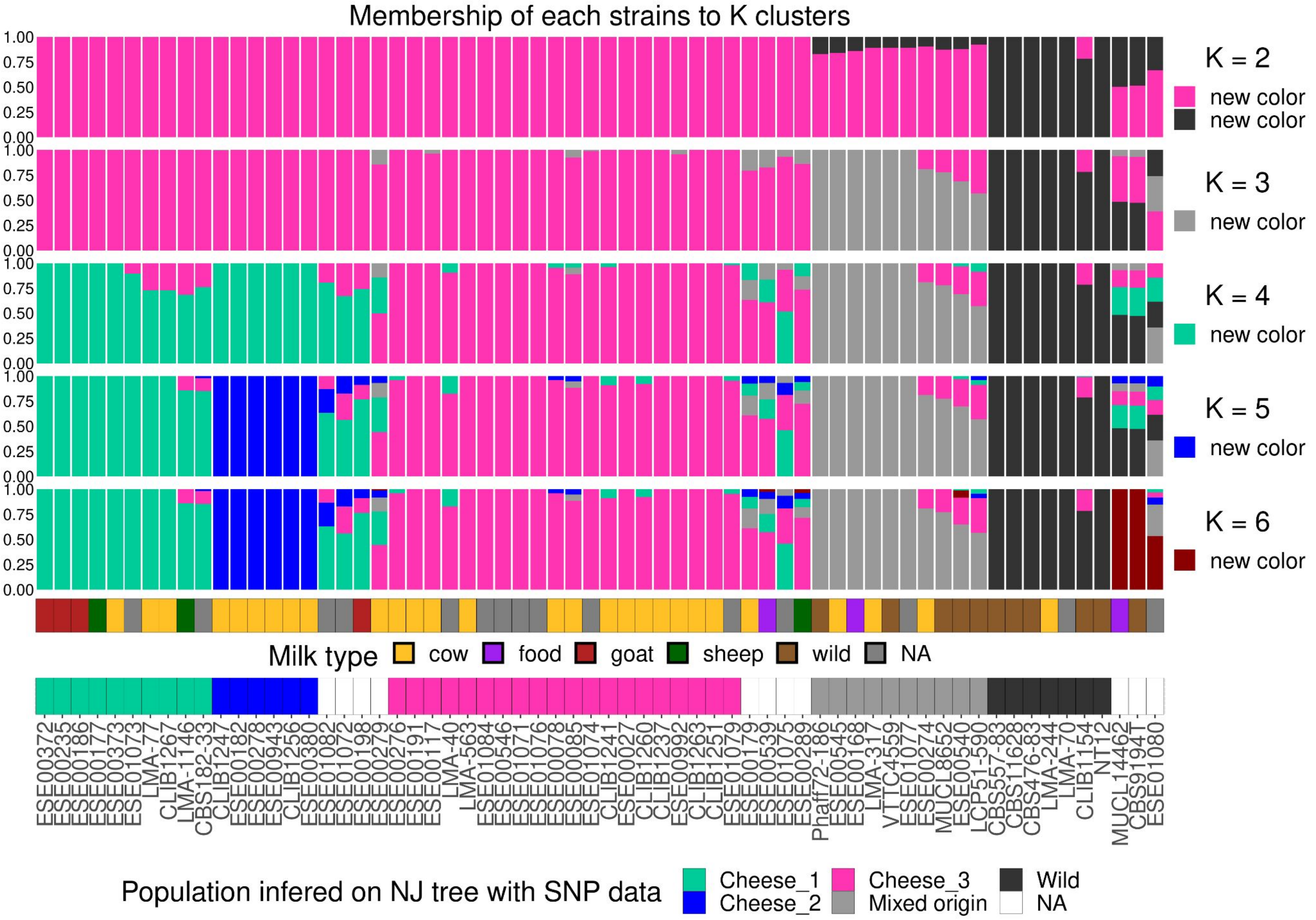


Figure S1: Population structure of *Geotrichum candidum*.

Population subdivision inferred for *K* populations ranging from two to six. Colored bars represent the coefficients of membership in the K gene pools based on genomic data. Each bar represents a strain, its name being indicated at the bottom of the figure. The new color for each *K* increment is indicated on the right part. The second order rate of change in the likelihood (ΔK) peaked at *K*=6. The names of the five populations considered in experiments are indicated on the last rows. Below the admixture plot, two rows indicate the type of milk from which the strains were sampled and the population subdivision.

Figure S2

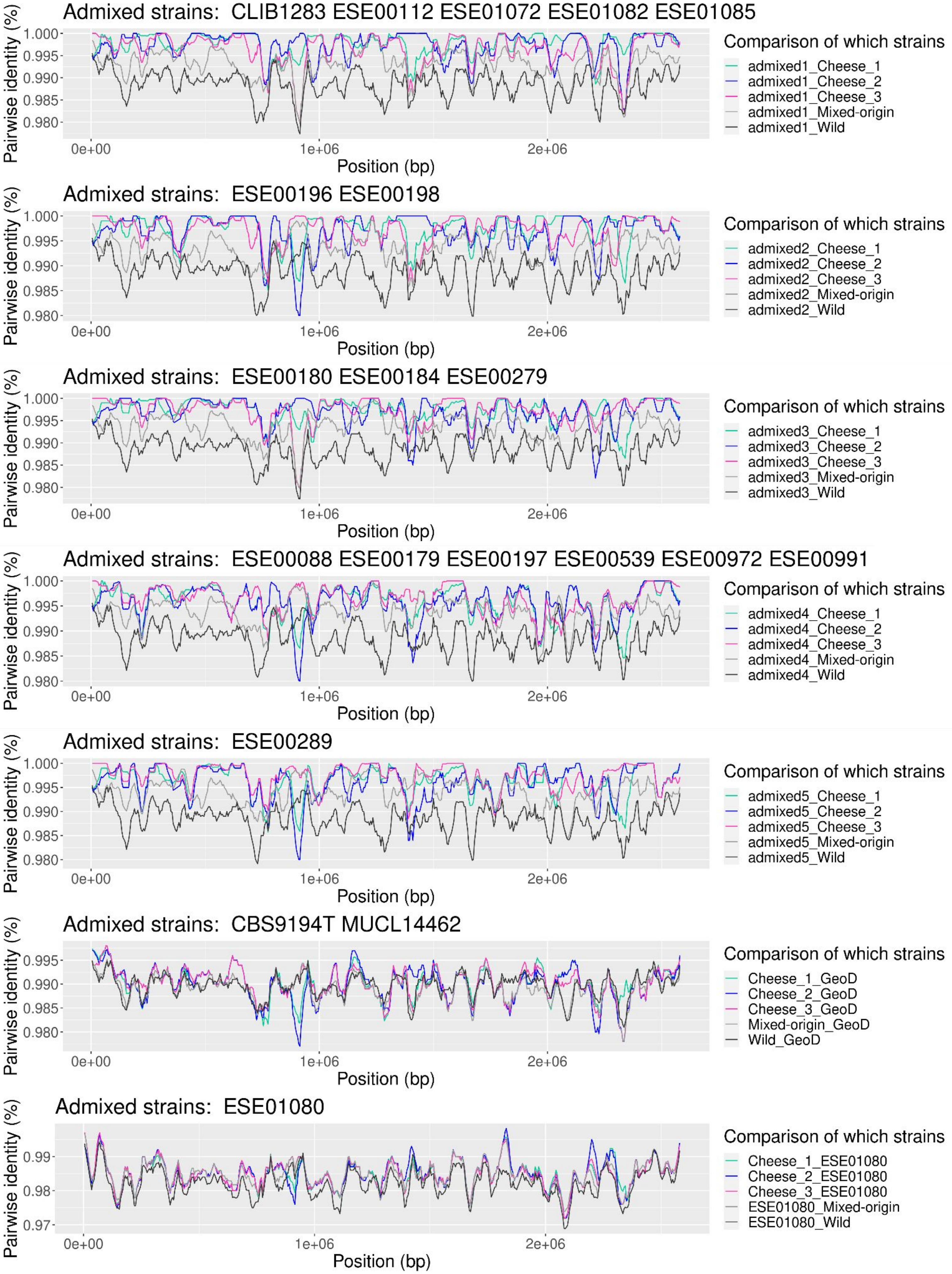
Admixed strains: CLIB1283 ESE00112 ESE01072 ESE01082 ESE01085
Comparison of which strains
admixed1_Cheese_1
admixed1_Cheese_2
admixed1_Cheese_3
admixed1_Mixed-origin
admixed1_Wild
Admixed strains: ESE00196 ESE00198
Comparison of which strains
admixed2_Cheese_1
admixed2_Cheese_2
admixed2_Cheese_3
admixed2_Mixed-origin
admixed2_Wild
Admixed strains: ESE00180 ESE00184 ESE00279
Comparison of which strains
admixed3_Cheese_1
admixed3_Cheese_2
admixed3_Cheese_3
admixed3_Mixed-origin
admixed3_Wild
Admixed strains: ESE00088 ESE00179 ESE00197 ESE00539 ESE00972 ESE00991
Comparison of which strains
admixed4_Cheese_1
admixed4_Cheese_2
admixed4_Cheese_3
admixed4_Mixed-origin
admixed4_Wild
Admixed strains: ESE00289
Comparison of which strains
admixed5_Cheese_1
admixed5_Cheese_2
admixed5_Cheese_3
admixed5_Mixed-origin
admixed5_Wild
Admixed strains: CBS9194T MUCL14462
Comparison of which strains
Cheese_1_GeoD
Cheese_2_GeoD
Cheese_3_GeoD
Mixed-origin_GeoD
Wild_GeoD
Admixed strains: ESE01080
Comparison of which strains
Cheese_1_ESE01080
Cheese_2_ESE01080
Cheese_3_ESE01080
ESE01080_Mixed-origin
ESE01080_Wild
Pairwise identity (%)
Position (bp)
0e+00
1e+06
2e+06

Figure S2: Pairwise identity between admixed and other strains, averaged per population of *Geotrichum candidum.*

Pairwise identity along the genome, averaged per population, with only the first scaffold of the CLIB 918 genome shown

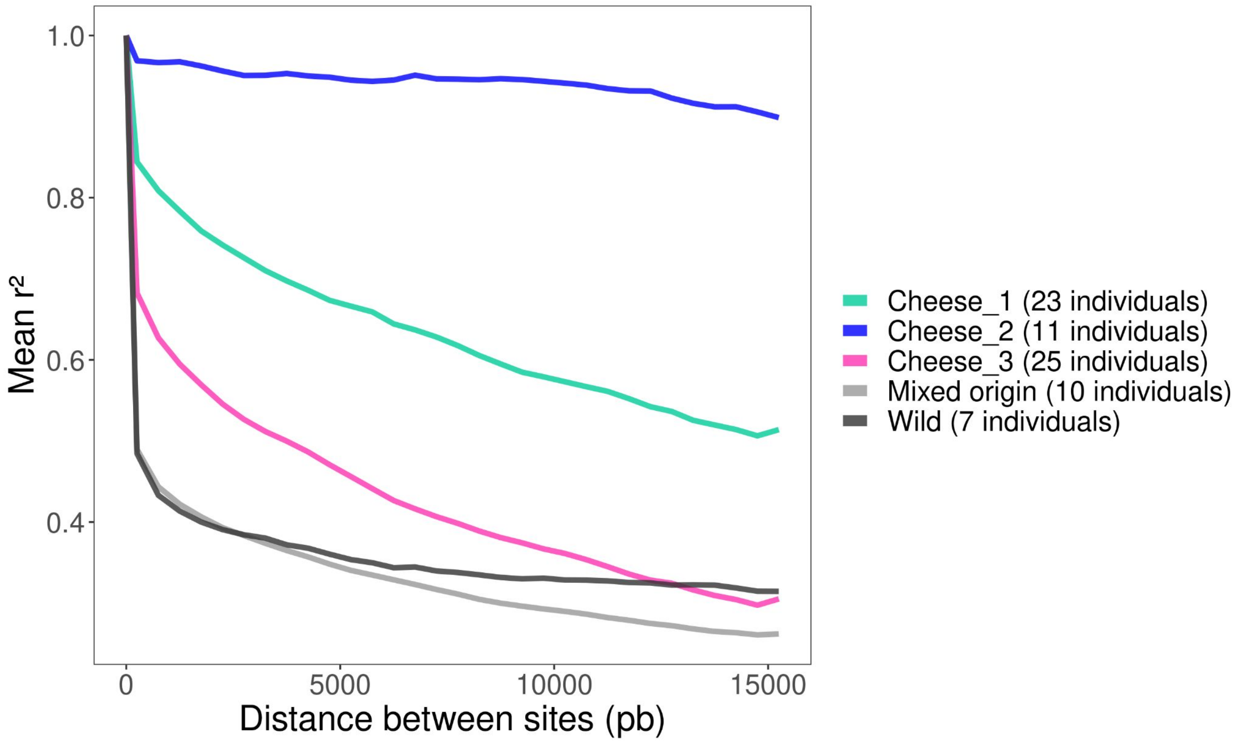


Figure S3: Linkage disequilibrium against distance between SNPs for the five *Geotrichum candidum* populations.

The $r^2$ is shown, representing the square of the correlation coefficient between two indicator variables; $r^2$ varies between 0 when two markers are associated at random and 1 when they provide identical information.

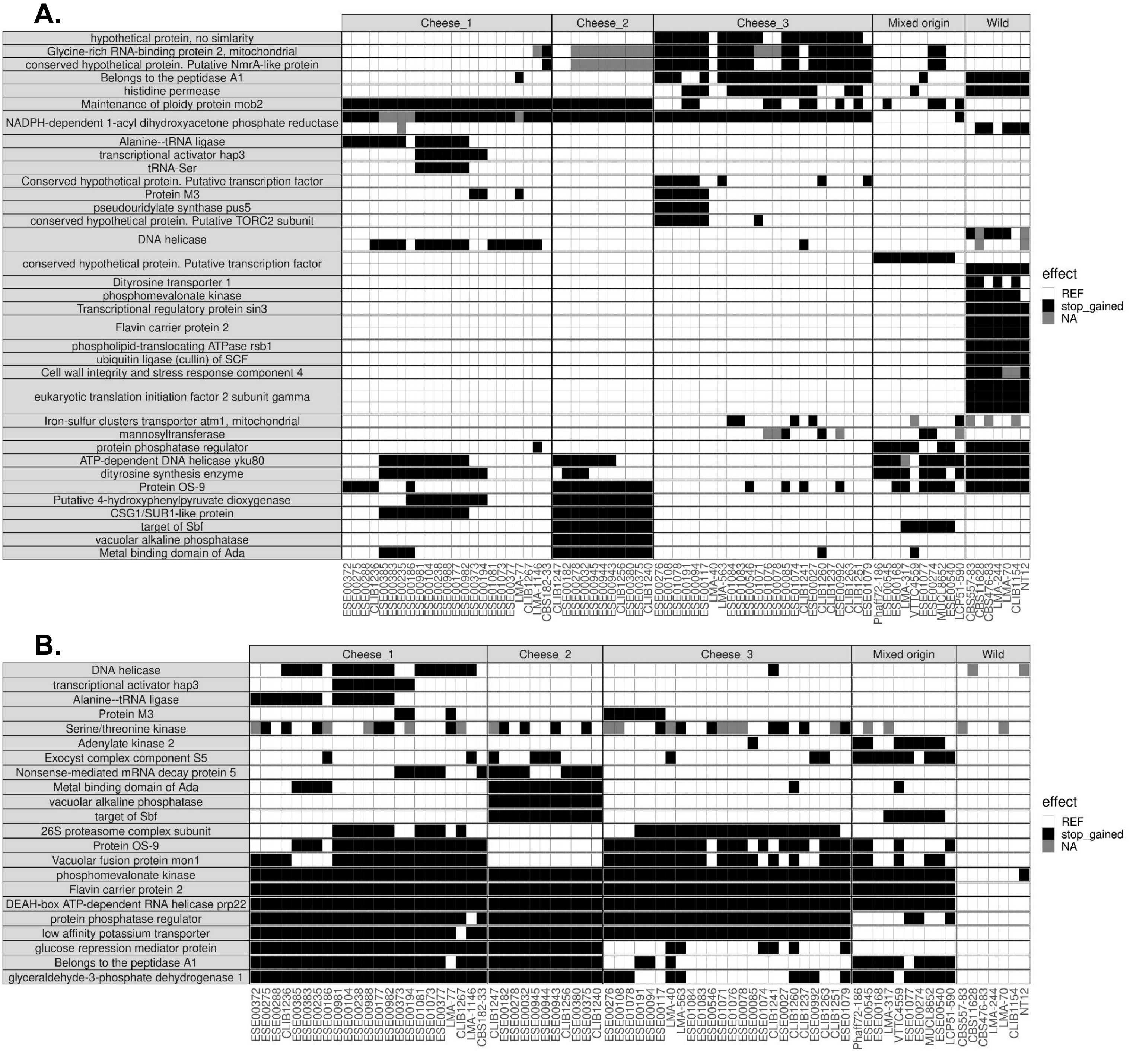


Figure S4: Single nucleotide polymorphism (SNPs) inducing a premature stop codon for each *Geotrichum candidum* strain.

We only show Stop -inducing SNPs present in more than three strains. Columns represent strains, ordered according to the maximum likelihood (ML) tree. Cells are colored in black when the corresponding SNP induces a premature stop codon, white when there is no substitution for this site compared to the reference genome and grey when the SNP status could not be assessed, substitution that induces other effects on the protein sequence were not present in this subset of Stop-inducing sites. Each row is a site; when multiple SNPs induced stop codons in the same gene, the corresponding rows were grouped and separated from other genes by black lines. The analysis was done using either the CLIB 918 (Cheese_3) genome (A) or the LMA-244 (Wild) genome (B) as a reference.

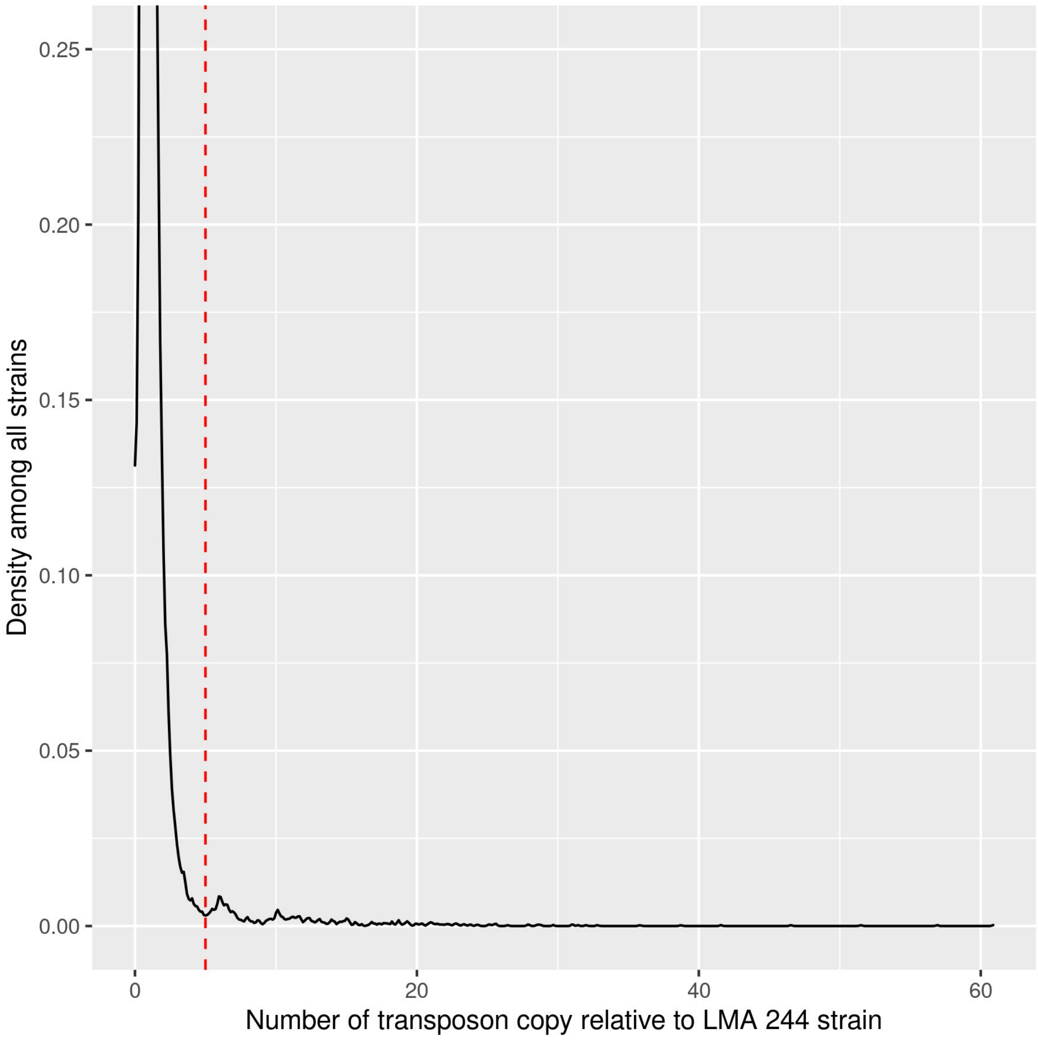


Figure S5: Density of transposable element copy number relative to the LMA-244 strain

To better show the fat tail distribution, the y-axis (density) was cut at 25%. A red dashed line indicates the threshold of five times more copy number as the LMA-244 strain.

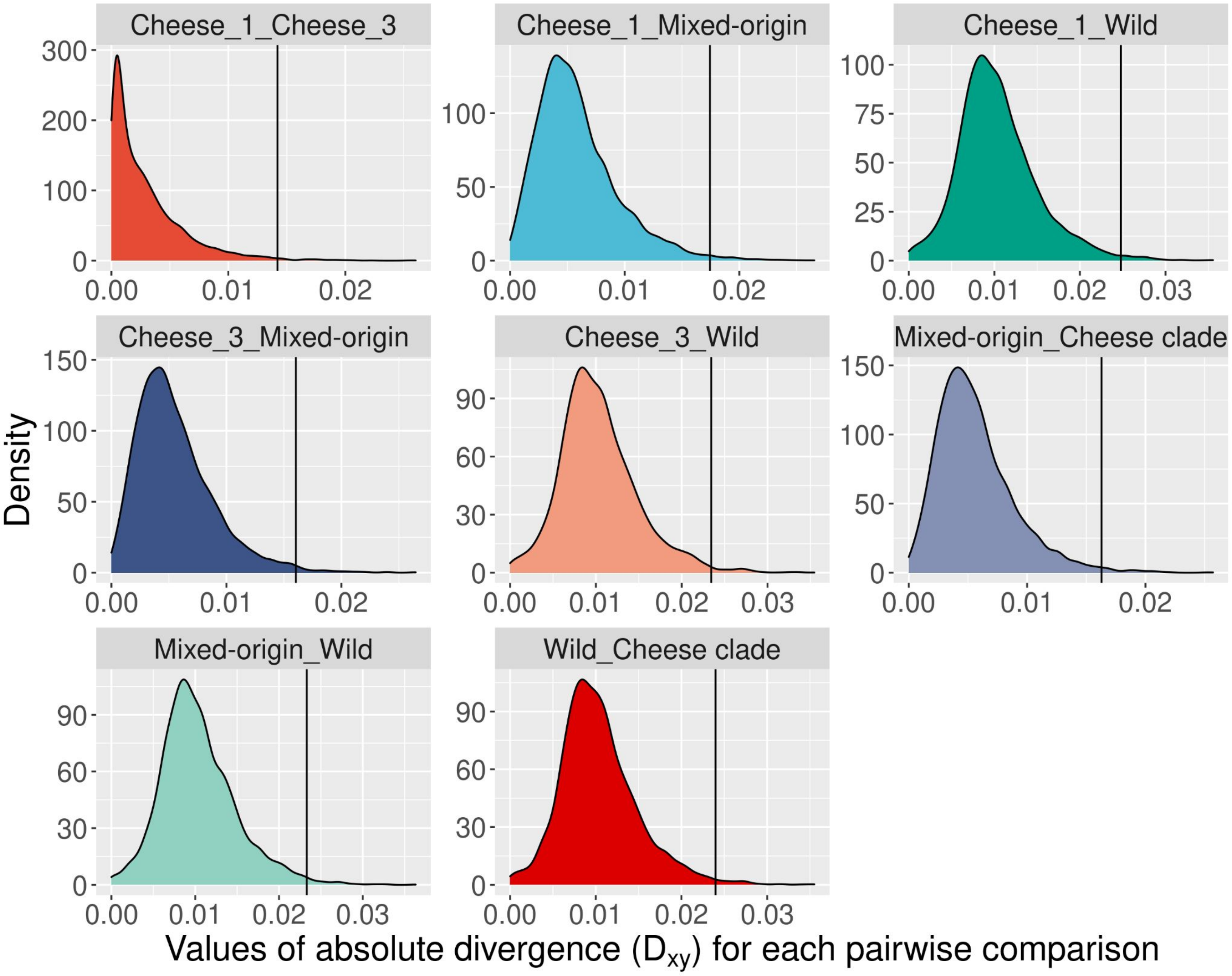


Figure S6: Distributions of absolute divergence ($d_{xy}$) values for different pairs of populations.

Distribution of absolute divergence ($d_{xy}$) values for each pairwise population comparison from the genomic scan analysis. Density is given as an overlapping window number for a specific value of $d_{xy}$, each window being 7.5 kb wide with a step of 5 kb (optimal values based on variants densities). A black vertical line indicates the threshold of 1% highest values kept for the enrichment test.

Figure S7

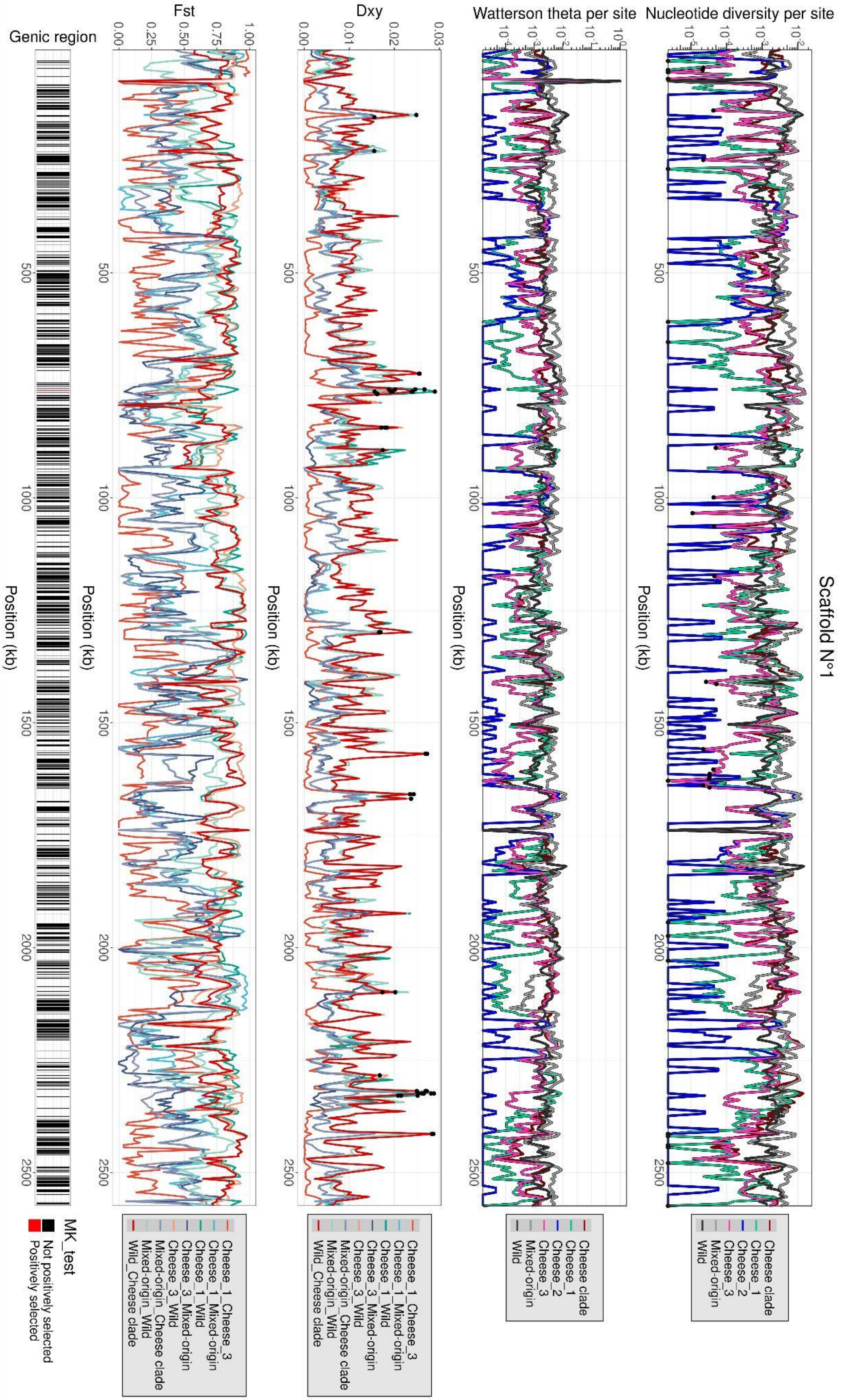

Scaffold N°1
Nucleotide diversity per site
Watterson theta per site
Dxy
Fst
Genic region
Position (kb)
Cheese clade
Cheese_1
Cheese_2
Cheese_3
Mixed-origin
Wild
Cheese_1_Cheese_3
Cheese_1_Mixed-origin
Cheese_1_Wild
Cheese_3_Mixed-origin
Cheese_3_Wild
Mixed-origin_Cheese clade
Mixed-origin_Wild
Wild_Cheese clade
MK_test
Not positively selected
Positively selected

Figure S7: Genomic scan of within-population genetic diversity and between-population differentiation in *Geotrichum candidum*.

Genomic of the nucleotide diversity $\pi$, watterson's theta $\theta_w$, absolute divergence $d_{xy}$ and fixation index $F_{ST}$ (Hudson et al., 1992) along the scaffold 1 of the CLIB 918 reference genome. At the bottom, a guide indicates genic regions in black and non genic regions in white. On the bottom of the figure, genic regions are shown in black (not positively selected) or red (positively selected in the McDonald and Kreitman test) rectangles. On the first panel (nucleotide diversity π), 5% lowest π values in the three cheese populations were highlighted by black dots. On the third panel (absolute divergence $d_{xy}$), the 1% highest values of $d_{xy}$ of each pairwise comparison are highlighted by back dots.

Figure S8

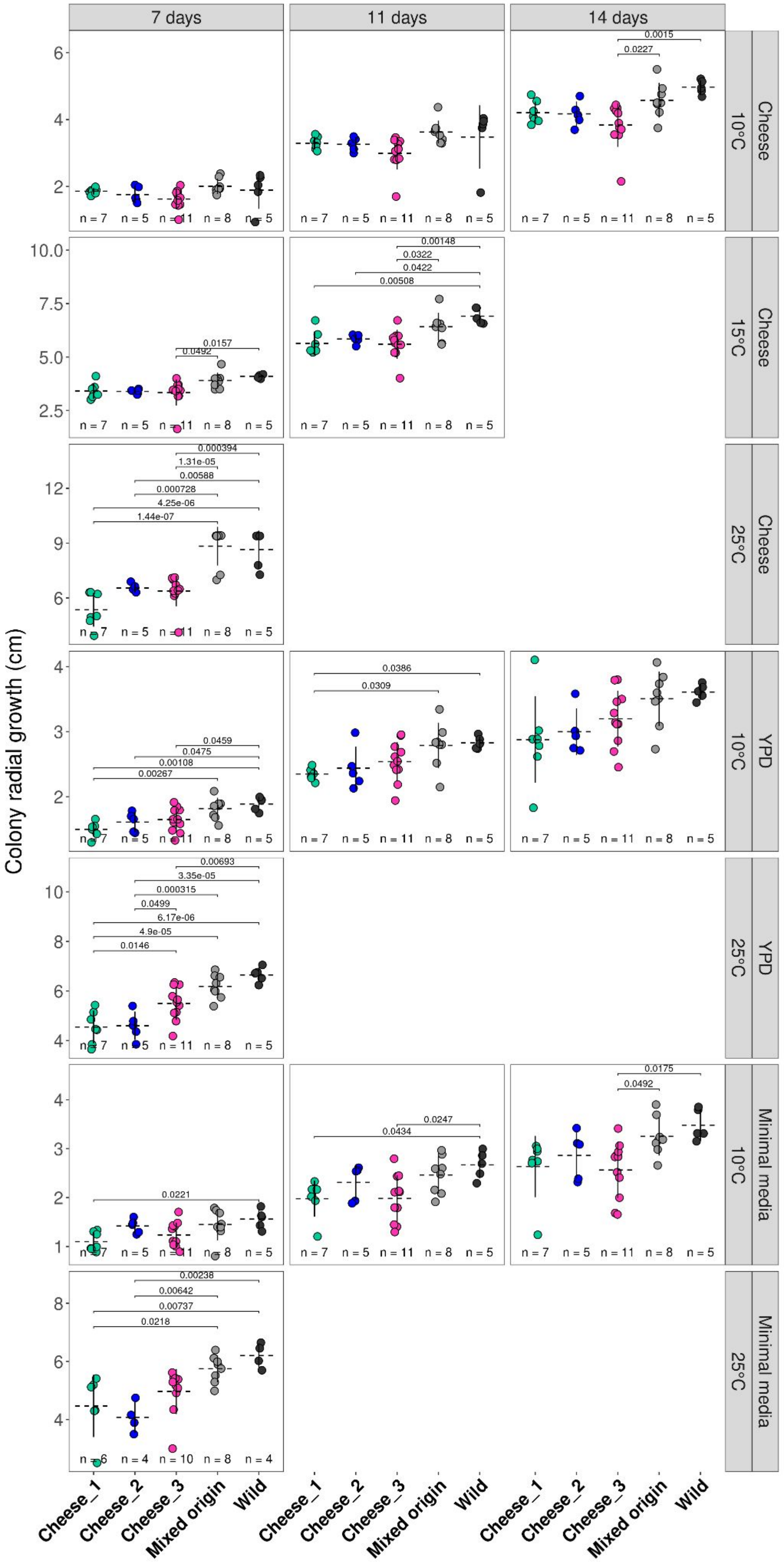

7 days
11 days
14 days
Cheese
10°C
15°C
25°C
YPD
Minimal media
Colony radial growth (cm)
Cheese_1
Cheese_2
Cheese_3
Mixed origin
Wild

Figure S8: Differences in growth between the five populations of *Geotrichum candidum* populations for different media and temperatures

Mean radial growth of the three cheese populations, mixed-origin and wild populations on cheese (1% salt), yeast peptone dextrose (YPD) and minimal media at 10, 15 and 25°C for 7, 11 and 14 days.

Each point represents a strain, horizontal dotted lines and vertical lines represent the mean and the standard deviation of the population, respectively. The number *n* at the bottom of plots indicates the number of strains per population used for measuring these phenotypes. Significant pairwise Tukey tests are indicated with brackets and p-values.

Figure S9

**A.**

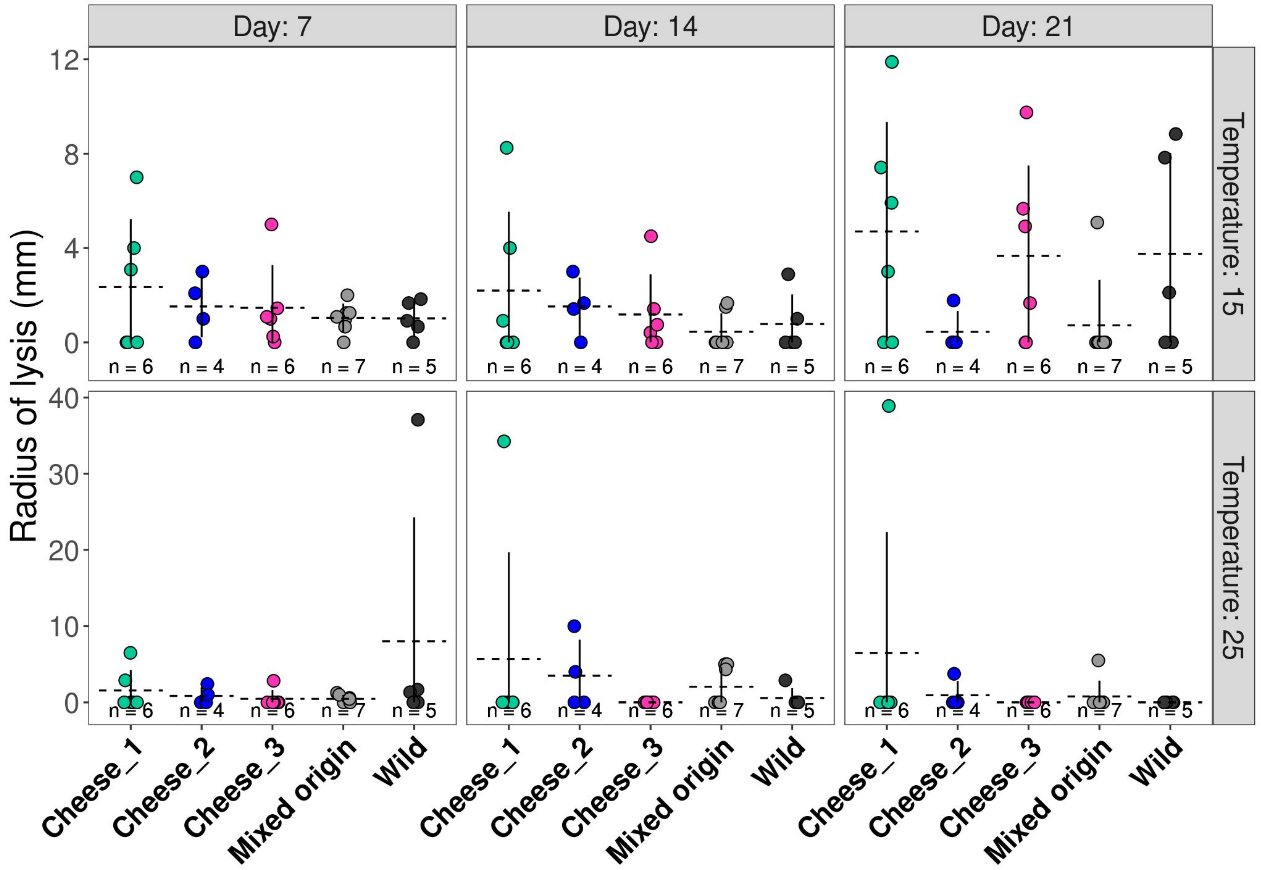

Day: 7
Day: 14
Day: 21
Temperature: 15
Temperature: 25
Radius of lysis (mm)
n = 6
n = 4
n = 6
n = 7
n = 5
Cheese_1
Cheese_2
Cheese_3
Mixed origin
Wild


**B.**

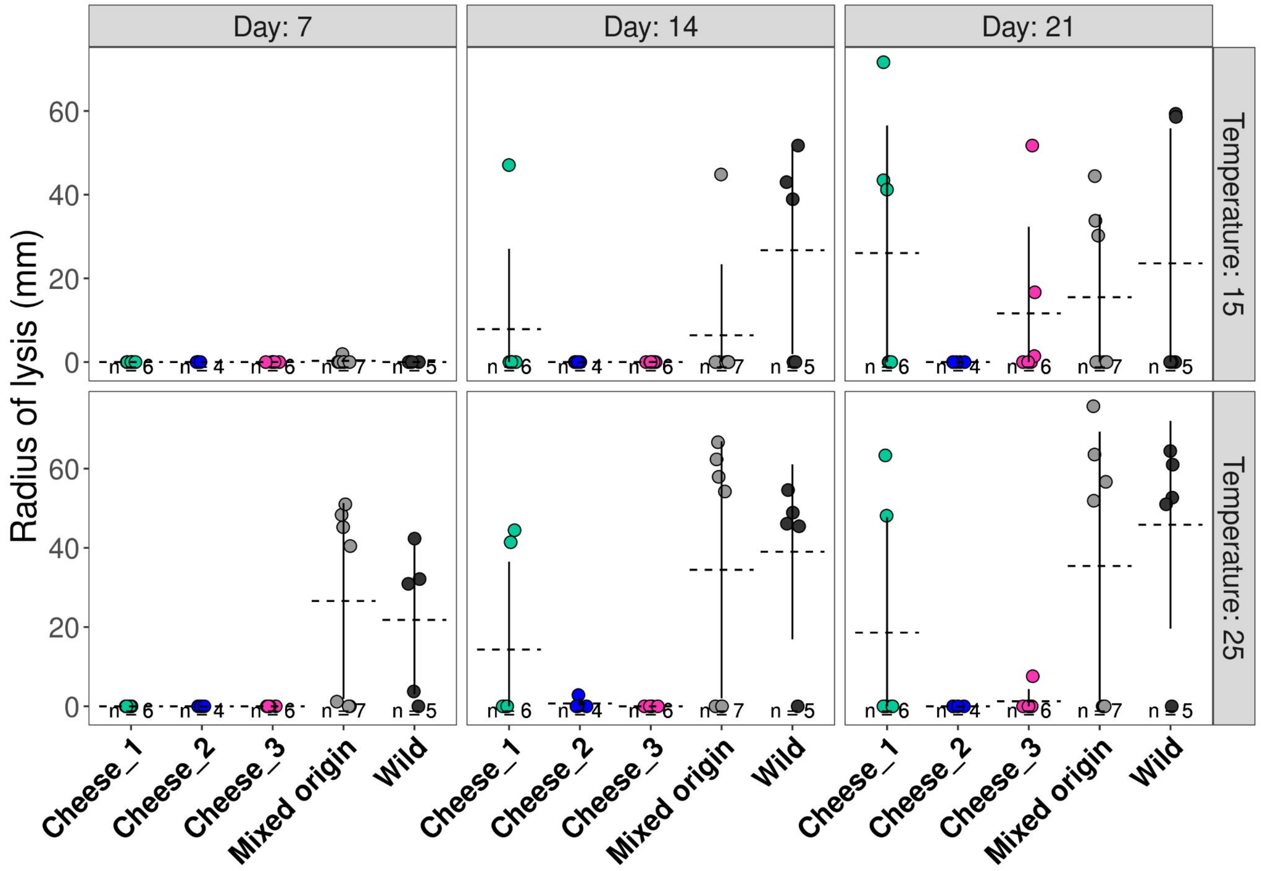

Day: 7
Day: 14
Day: 21
Temperature: 15
Temperature: 25
Radius of lysis (mm)
n = 6
n = 4
n = 6
n = 7
n = 5
Cheese_1
Cheese_2
Cheese_3
Mixed origin
Wild

Figure S9: Differences in lipolytic and proteolytic activity among the five populations of *Geotrichum candidum* populations for different growing time and temperature

A: Lipolytic activity of *G. candidum* at 15 and 25°C and grown for 7, 14 and 21 days.

B: Proteolytic activity of *G. candidum* at 15 and 25°C and grown for 7, 14 and 21 days.

Each point represents a strain, horizontal dotted lines and vertical lines represent the mean and the standard deviation of the population respectively. The number *n* at the bottom of plots indicates the number of strains per population used for measuring these phenotypes. Significant pairwise Tukey tests are indicated with brackets and p-values.

Figure S10

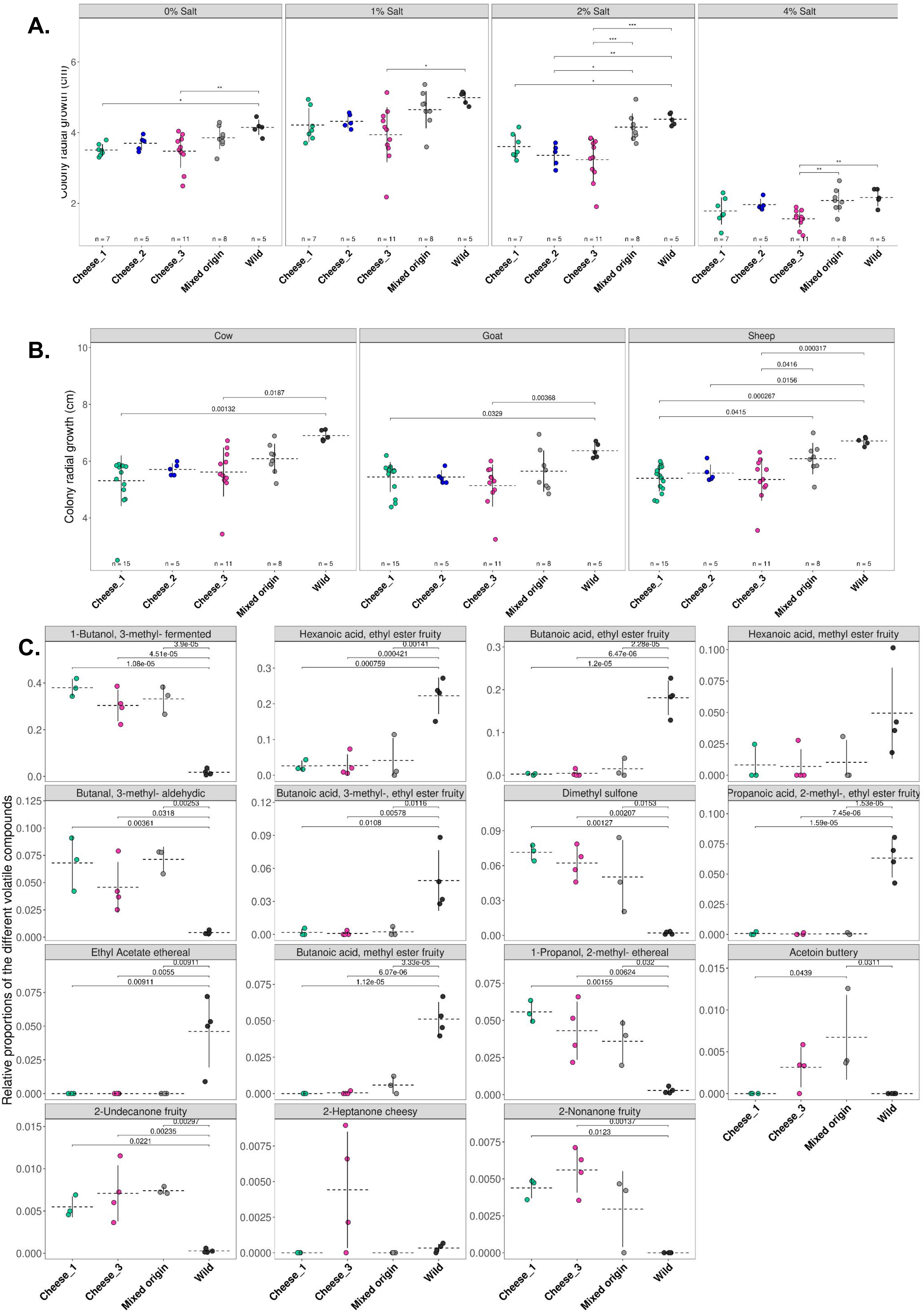

A.
0% Salt
1% Salt
2% Salt
4% Salt
Colony radial growth (cm)
n = 7
n = 5
n = 11
n = 8
n = 5
Cheese_1
Cheese_2
Cheese_3
Mixed origin
Wild
B.
Cow
Goat
Sheep
Colony radial growth (cm)
0.0187
0.00132
0.00368
0.0329
0.000317
0.0416
0.0156
0.000267
0.0415
n = 15
n = 5
n = 11
n = 8
n = 5
C.
Relative proportions of the different volatile compounds
1-Butanol, 3-methyl- fermented
3.9e-05
4.51e-05
1.08e-05
Hexanoic acid, ethyl ester fruity
0.00141
0.000421
0.000759
Butanoic acid, ethyl ester fruity
2.28e-05
6.47e-06
1.2e-05
Hexanoic acid, methyl ester fruity
Butanal, 3-methyl- aldehydic
0.00253
0.0318
0.00361
Butanoic acid, 3-methyl-, ethyl ester fruity
0.0116
0.00578
0.0108
Dimethyl sulfone
0.0153
0.00207
0.00127
Propanoic acid, 2-methyl-, ethyl ester fruity
1.53e-05
7.45e-06
1.59e-05
Ethyl Acetate ethereal
0.00911
0.0055
0.00911
Butanoic acid, methyl ester fruity
3.33e-05
6.07e-06
1.12e-05
1-Propanol, 2-methyl- ethereal
0.032
0.00624
0.00155
Acetoin buttery
0.0311
0.0439
2-Undecanone fruity
0.00297
0.00235
0.0221
2-Heptanone cheesy
2-Nonanone fruity
0.00137
0.0123
Cheese_1
Cheese_3
Mixed origin
Wild

Figure S10: Differences in salt tolerance, growth on different milk types and volatile compounds between the five populations of *Geotrichum candidum.*

Each point represents a strain, horizontal dotted lines and vertical lines represent the mean and the standard deviation of the phenotype in the population, respectively. The number *n* at the bottom of plots indicates the number of strains used per population for measuring the corresponding phenotypes. The pairwise Tukey tests performed to assess whether there were mean differences between populations are indicated with brackets and their p-values are given.
(A) Mean radial growth at 10°C of the three cheese, mixed-origin and wild populations of *G. candidum* on cheese agar medium with different salt concentrations: unsalted, 1% salt for mimicking St Nectaire and cream cheeses, 2% salt for mimicking Camembert and goat cheeses and 4% salt for mimicking blue cheeses.
(B) Mean radial growth at 10°C of the three cheese populations, the mixed-origin and the wild populations on yogurt agar media made with raw cow, goat and sheep milks.
(C) Relative proportions of major volatile compounds in Cheese_1, Cheese_3, mixed-origin and wild populations of *G. candidum.* The volatile compounds shown were those contributing the most to the two first PCA axes or those that are important for cheese ripening. For each compound, the related corresponding descriptor from thegoodscentscompany.com was added.
