## Supplementary Table legend for "Domestication of different varieties in the cheese-making fungus *Geotrichum candidum*"

Table S5: Population genetics statistics estimating genetic diversity ($\pi$, watterson's θ and Tajima's D) in the five *Geotrichum candidum* populations and populations of three other fungal species (*Penicillium roqueforti, Penicillium camemberti* and *Saccharomyces cerevisiae*) (Dumas et al., 2020; Peter et al., 2018; Ropars et al., 2020b).

Cells are colored from the lowest (white) and the highest (red) value of each indices.

Table S6: Distribution of the two mating types in each population and test of the deviation from a 1:1 ratio.

Table S7: Phi test of each population of *Geotrichum candidum* using the first scaffold of CLIB 918 assembly.

The pairwise homoplasy index (PHI) test assesses with permutations the null hypothesis of no recombination by looking at the genealogical association among adjacent sites.

Table S8: Number and percentage of single nucleotide polymorphism (SNPs) classified by impact, functional class, effect and genomic regions for each *Geotrichum candidum* population.

Based on the 7,150 coding sequences (CDS) of the CLIB 918 assembly, the effect of all variants was assessed. Each variant is categorized in different functional effects on the protein sequence. Results are averaged per population and shown either as the mean number or percentage of SNPs. Results were computed using snpeff (Cingolani et al., 2012).

(A) Total number of single nucleotide polymorphisms (SNPs) within each *G. candidum* population.

(B) Putative variant impact prediction. Different impacts categories are defined in Snpeff manual in 'Input & output files' section

(C) Protein sequence effect of SNPs. Variants can either not change amino acid sequence (silent), change the amino acid (missense) or induce stop codons (nonsense)

(D) Functional effect of SNPs defined in Snpeff manual in 'Input & output files' section

(E) Position of the SNPs compared to genes

Table S9: Copy number variants differentiating G*eotrichum candidum* populations and test for enrichment in gene ontologies contained within these windows.

(A) Number of windows that differentiate clades based on copy number variation using LMA 244 assembly as reference

(B) Number of windows that differentiate clades based on copy number variation using CLIB 918 assembly as reference

(C) Enrichment test on gene ontologies (GO) present within the windows that differentiate clades based on copy number variation using CLIB 918 assembly as reference

(D) Enrichment test on gene ontologies (GO) present within the windows that differentiate clades based on copy number variation using LMA 244 assembly as reference

Table S10: Repeat copy number for the different strains of *Geotrichum candidum.*
